# Cross-feeding options define eco-evolutionary dynamics of deep oligotrophic groundwater microbiome

**DOI:** 10.1101/2024.08.02.606368

**Authors:** Maryam Rezaei Somee, Carolina González-Rosales, Matti Gralka, Stephanie Turner, Stefan Bertilsson, Mark Dopson, Maliheh Mehrshad

## Abstract

Deep groundwaters populated by diverse and active microbes are among the most energy and nutrient-limited ecosystems. Characteristics of this ecosystem including nutrient and dispersal limitations, low cell densities, and episodic growth strategy interactively underpin the so far elusive eco-evolutionary dynamics of its microbiome. Here we applied a modular metabolic analyses on genome-resolved reconstructed community of disconnected deep groundwaters in the Fennoscandian Shield. In the community of deeper groundwaters despite their highly oligotrophic nature, lineages with larger genomes maintained larger populations which we hypothesize to be connected with their limited cross-feeding options. Thus providing an extension on the streamlining theory emphasizing the importance of ecological interactions in genome evolution which is further supported by the observed decrease in abundance of lineages with known metabolic dependencies, such as Patescibacteria and DPANN, with depth. The modular metabolic analyses showed that remarkably common niches based on same cross-feeding interactions are also available in different groundwaters, in addition to common niches for primary production. While these shared niches are critical for community assembly in this ecosystem, in different boreholes different lineages populated them. Our results provided new insights into the role of metabolic cross-feeding in genome evolution and community assembly of deep groundwater microbiome.

---

Deep oligotrophic groundwaters are among the most energy and nutrient-limited ecosystems on our planet. Yet, they sustain diverse and metabolically active communities^1,2^ that encompass representatives from all domains of life as well as viruses^3,4^. It has been shown that available nutrients, reducing agents, and geochemical features of the bedrock hosting groundwaters^5^ define the community composition of deep groundwater microbes. Recent studies also reveal the existence of a common “core” microbiome in Fennoscandian Shield deep groundwaters^3^ that is mainly shaped by ecological convergence of species in communities inhabiting similar geologies^3^. Beyond the importance of niche dimensions (both biotic and abiotic) for shaping the composition of deep groundwater communities^3^, it is not fully understood how the characteristic features of this milieu affect the eco-evolutionary dynamics of its microbiota.

Despite ongoing debate around the role of environmental factors in the evolution of genome features (i.e., genome size and GC content)^6^, the assumption that genomes with lower GC content and smaller genome size can be more abundant in nutrient-limited ecosystems^7^ seems to have been confirmed at least for oligotrophic niches in the surface ocean and lakes^8–10^. Abundant lineages with streamlined genomes that dominate these habitats are hypothesized to compensate for auxotrophies and metabolic dependencies via cross-feeding of externally supplied substrates or from tight symbiotic interactions with other community members^11^, putting the black queen hypothesis in play for penalizing further loss of essential functions at the level of communities^12^. Following the same logic, the lower energy and nutrient demands of lineages with smaller genome sizes and lower GC content should in theory allow them to maintain larger populations in the prevailing oligotrophic conditions of the deep groundwaters. However, fundamental differences in dispersal methods and symbiosis strategies of deep groundwater microbiome together with limited nutrient access and poor metabolic cross-feeding options due to cell sparsity^11^ as well as their episodic growth strategies^3^, could potentially contribute to the emergence of different eco-evolutionary avenues that remain elusive.

Deep groundwater ecosystems are disconnected from inputs of labile organic matter from the solar- exposed surface of our planet, and compounds able to reach deep niches (e.g., terrigenous lignin) are barely degradable under the prevailing anaerobic conditions^13^. Thus, conducive carbon acquisition strategies in deep groundwaters are scarce and limited to carbon fixation and recycling of cell debris^14^. Additionally, the main available nitrogen sources are expected to be ammonium (originating from cell necrosis), nitrate (also used as a final electron acceptor), and N2^15^. Previous studies have suggested that deep oligotrophic groundwater microbes employ an episodic growth strategy to ensure their subsistence^3^. The residual impact of this episodic growth could affect their symbiosis and interdependencies due to interruptions in different metabolic pathways during each growth episode. Consequently, to understand metabolic interactions of the deep groundwater microbiome in the context of predominantly episodic growth, the abundance of specific modules in carbon and nitrogen acquisition pathways needs to be examined for the production of intermediate compounds potentially relevant for metabolic interactions within the community.

In this study, an extensive multi-omics dataset termed the “Fennoscandian Shield Genomic Database” (FSGD)^3^, has been supplemented with additional sequencing data from deep oligotrophic groundwaters of the Äspö Hard Rock Laboratory (Äspö HRL) in Sweden and Olkiluoto Island, Finland. Inspecting the genomic features of the microbial community inhabiting each borehole and the metabolic cross-feeding strategies they adopt for subsistence in the deep groundwater ecosystems provide novel clues towards the decisive role of metabolic cross-feeding in the eco-evolutionary dynamics of these microbes and add to the existing understanding of the streamlining theory.

## Results and discussion

### Genome size and GC content distribution varies in different boreholes

The FSGD contains 1876 metagenome assembled genomes (MAGs) and 114 single cell amplified genomes (SAGs) with ≥50% completeness and ≤5% contamination. These genomes originated from 43 metagenomes and 114 single cell amplified genomes (SAGs), including datasets detailed in Mehrshad et al.^3^ plus nine newly sequenced metagenomes (**Supplementary Table S1**). Clustering MAGs/SAGs at 95% average nucleotide identity yielded 1185 representative genome clusters, which were affiliated to 83 phyla and 153 classes (**Supplementary Table S2)**. The MAGs/SAGs GC content ranged from 25 to 73% and their completeness-corrected estimated genome size (EGS) was in the range of 0.66 to 10.34 MB, with the majority of MAGs/SAGs having an EGS in the range of 1.25 to 2.5 Mb (**Fig. 1a**). An overall correlation between the EGS and GC content was also detected for the FSGD MAGs/SAGs (**Fig. 1a**). The distribution of MAGs/SAGs along the GC content range had a major peak at around 40% and two smaller peaks at around 57 and 63% (**Fig. 1a**). Nevertheless, MAGs/SAGs present in each disconnected borehole showed a different pattern with regards to the overall GC and EGS distribution (**Supplementary Figs. S1 and S2**). One borehole with strikingly different distribution compared to the overall pattern was the OL-KR46 (Olkiluoto Island, Finland) with MAGs/SAGs present in this borehole (*n*=27) featuring a GC peak at around 63% (**Supplementary Fig. S1**) and an EGS peak at around 3.7 Mb (**Supplementary Fig. S2**). Interestingly, the OL-KR46 borehole intersected the deepest FSGD groundwater at 528.7–531.5 meters below surface level (mbsl) and had the lowest Shannon index for alpha diversity (**Fig. 1b**). The measured salinity of 18 parts per thousand (ppt) for this borehole hints at an extended residence time of water in this borehole with persistent long-term disconnect from nutrient-rich surface waters^16^. Accumulation of genomes with larger EGS and higher GC content in this extremely nutrient-limited habitat showed that there was a higher cost to benefit for streamlining in this ecosystem as compared to the higher abundance of streamlined genomes in oligotrophic ocean and lake ecosystems^8^.

**Figure 1:**
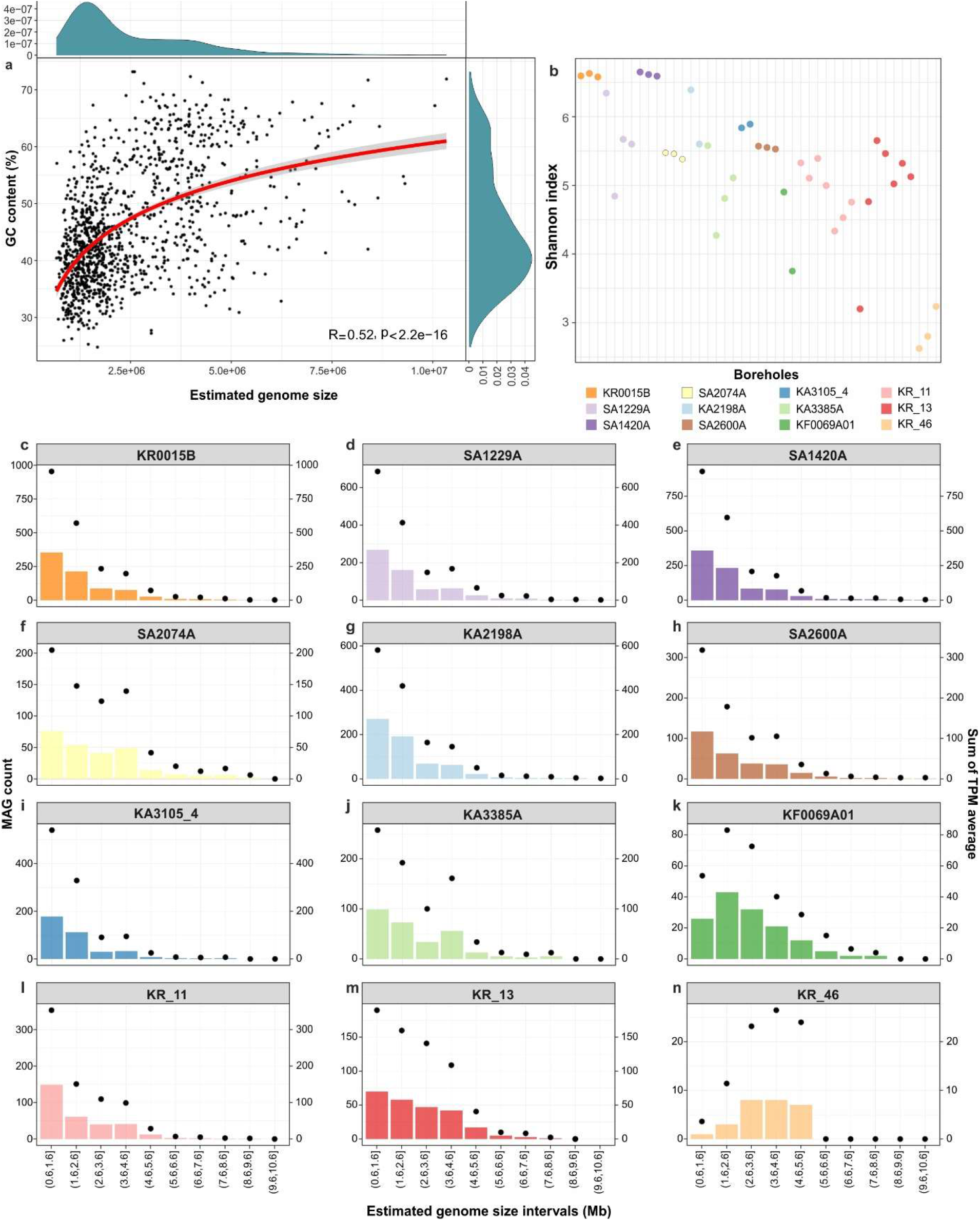
Distribution and prevalence of FSGD MAGs/SAGs in different deep groundwater boreholes. a) Correlation between GC content and estimated genome size (EGS) of all representative MAGs/SAGs. Side graphs illustrate the distribution density of MAGs/SAGs across GC and EGS spectrum. b) Alpha diversity for each borehole calculated based on the Shannon-Wiener index for representative MAGs/SAGs present in each borehole (those with nonzero log10 value of the calculated transcript per million (TPM)). c-n) The number of FSGD MAGs/SAGs in 1 Mb intervals along the estimated genome size spectrum are shown as bar plots and are separated in different panels for different boreholes. The overall abundance of MAGs/SAGs in each interval (average of nonzero TPM values in all metagenomes sequenced for each borehole) are overlaid on each panel as dot plots.

The abundance (i.e., population sizes) of representative MAGs with different EGS (grouped in ten categories at 1Mb intervals) in different boreholes, is critical for deciphering eco-evolutionary dynamics of these microbes. Factoring in the abundance of microbes inhabiting each borehole showed that genomes with an EGS between 0.6 to 2.6 Mb had the highest cumulative abundance in all boreholes (**Fig. 1c-n**) except for KF0069A01 (depth 454.8 mbsl, salinity 24 ppt, two samples, and 143 MAGs) and OL-KR46 (depth 528.7–531.5, salinity 18 ppt, three samples, and 27 MAGs). For these two boreholes, the average of nonzero transcript per million (TPM) abundance value of their MAGs/SAGs showed that genomes in the size range of 3.6 to 4.6 Mb had the highest population size (**Fig. 1k&n**). MAGs/SAGs detected in the KF0069A01 and OL-KR46 boreholes with an EGS in the range of 0.6 to 1.6 Mb mainly belonged to Patescibacteria (11.2 and 3.7% of MAGs/SAGs present in the borehole, respectively) and DPANN (2.1 and 0% of MAGs/SAGs present in the borehole, respectively). The shift in the peak of genome size could be partially attributed to the decreasing prevalence of Patescibacteria and DPANN within the community of the deepest boreholes both at the Äspö HRL and the Olkiluoto locations (**Supplementary Fig. S3**). This decrease in prevalence could be a consequence of reduced symbiotic encounters in the deeper groundwater specifically since it is shown that symbiotic interactions are critical for survival of representatives of these lineages^17,18^.

A strong correlation between the portion of noncoding DNA in the genome and either genome size (**Supplementary Fig. S4)** or GC content (**Supplementary Fig. S5**), was only detected for boreholes OL- KR46 (R=-0.68, p=9e-05). This imply that microbes potentially minimize the cost of replication by reducing the noncoding fraction of their genomes. However, the percentage of their non-coding DNA does not drop below 9% which is still far from the 5% reported for streamlined SAR11^8^ in the ocean.

The population size of representative MAGs/SAGs within each borehole, calculated as an average of their non-zero abundances (log10 of TPM value) in all samples of that borehole, showed a normal distribution for most boreholes (**Supplementary Fig. S6**). There was no strong correlation between the calculated population size and EGS (**Supplementary Fig. S7**) or the GC content of MAGs/SAGs in each borehole (**Supplementary Fig. S8**). This highlighted that lineages from across the range of EGS and GC content can develop large populations. Further dividing reconstructed MAGs/SAGs into different phyla also detected a range of different population sizes among representatives of each phylum **(Supplementary Fig. S9)**.

### Carbon fixation is more common and efficient in high GC content lineages

To define the role of metabolic capabilities in eco-evolutionary dynamics of deep groundwater microbiomes, representative FSGD MAGs/SAGs were surveyed for the presence/absence of genes encoding carbon fixation and nitrogen acquisition pathways. To account for their episodic growth in response to episodic availability of nutrients^3^, a modular metabolic analysis was performed. For this approach C-fixation pathways (reductive citrate cycle (rTCA), 3-hydroxy-propionate bi-cycle (3HP), dicarboxylate-hydroxybutyrate, hydroxy-propionate-hydroxy-butylate (HPHB), Calvin-Benson-Bassham cycle (CBB), reductive acetyl-CoA/Wood-Ljungdahl pathway (WLP), phosphate acetyltransferase-acetate kinase (PAT-ACK)) were broken down to 58 modules based on KEGG (containing 168 genes; **Fig. 2**). These modules result in the production of intermediate compounds that could potentially be used for cross-feeding and therefore, 25 transporters (encoded by 39 genes) specialized for acquisition/export of these intermediates from the milieu were also surveyed. Furthermore, five modules involved in inorganic N-acquisition (ten genes encoding nitrogen fixation, nitrate/nitrite assimilation, nitrate/nitrite dissimilation plus ten additional genes encoding ammonium permease and NO3/NO2 transporters) were included (full list of modules and their enzymes in **Supplementary Table S3**).

**Figure 2:**
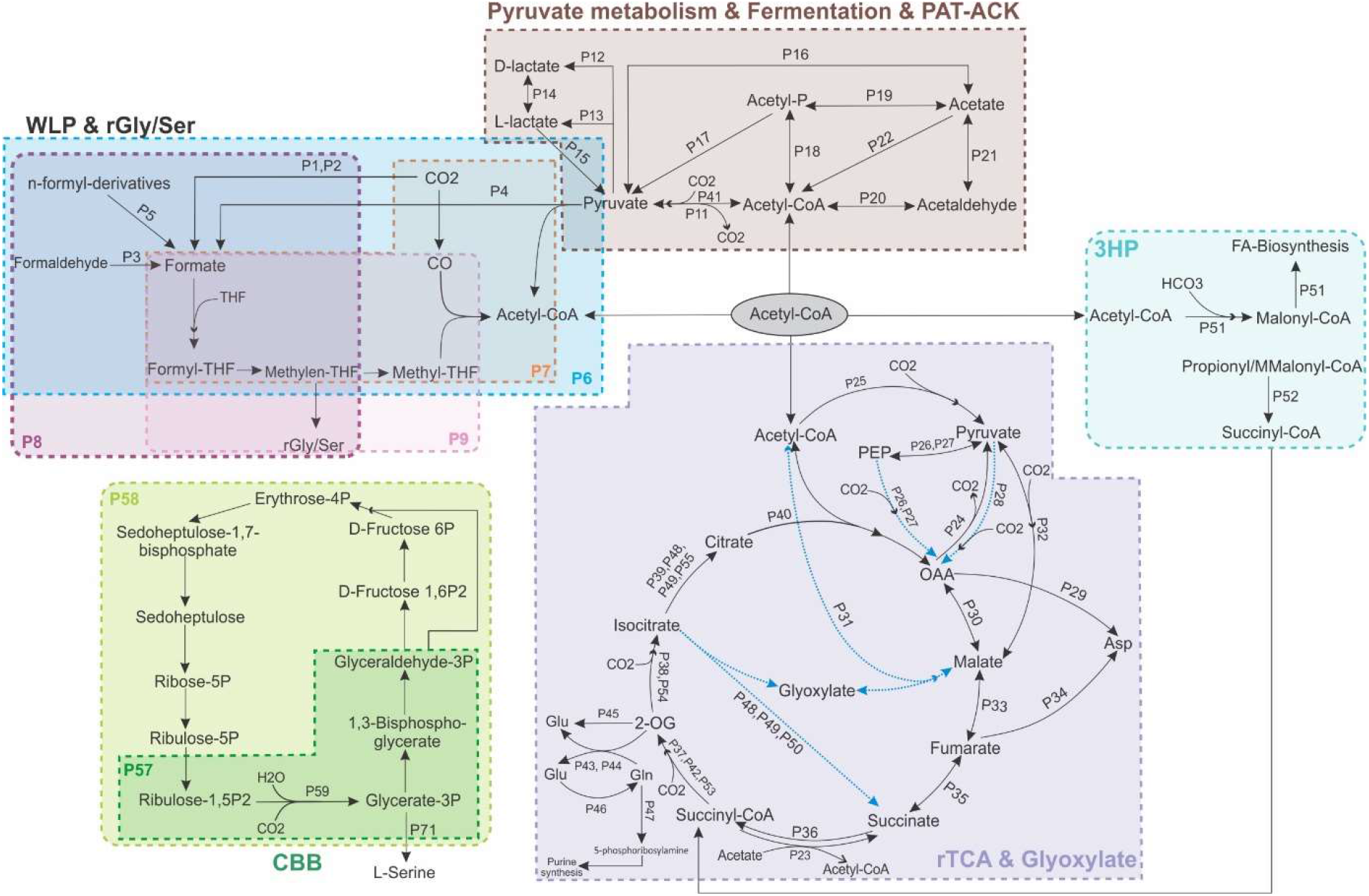
Schematic overview of carbon fixation pathways and designated metabolic modules used in this study. Anaplerotic pathways are shown with gray arrows. Modules related to transporters are not included. Only pathways that were detected in recovered FSGD MAGs/SAGs are represented. KEGG orthology (KOs) encoding the hydroxy-propionate-hydroxy-butyrate pathway (HPHB) were not detected in FSGD MAGs/SAGs and the pathway is not included. Abbreviations are as follows, PAT- ACK: phosphate acetyltransferase-acetate kinase; WLP & rGly/Ser: Wood-Ljungdahl pathway and reductive glycine/serine; CBB: Calvin-Benson-Bassham cycle, rTCA: reductive tricarboxylic acid/reductive citrate cycle; 3HP: 3-hydroxypropionate bi-cycle.

Genes encoding the 15 enzymes involved in carbon fixation (CO2 or HCO3) in rTCA, 3HP, HPHB, and WLP pathways were more prevalent among lineages with higher GC content and larger EGS except for the gene encoding CBB key enzyme Rubisco **(Supplementary Figs. S10&S11)**. The enzymes PEP carboxylase, PEP carboxy kinase (PEPCK), pyruvate carboxylase, crotonyl-CoA carboxylase/reductase, and formate dehydrogenase have a high affinity for capturing carbon under both saturation and low concentrations of CO2/HCO ^19^. These enzymes showed a higher prevalence among high GC-content FSGD lineages (**Supplementary Fig. S10**). In the deep groundwaters, the efficient capture of these main carbon sources by high GC content lineages indicated their significant contribution to carbon fixation.

WLP, reductive glycine (rGly), and rTCA fixation pathways are energy-efficient in having ATP requirement of less than 1, 1 (per acetyl-CoA), and 3 (per pyruvate), respectively^19,20^. Among these, the rTCA was more prevalent in high GC content lineages **(Supplementary Fig. S12)** and while the rTCA does not have the lowest ATP requirement, the anoxic deep groundwater is a conducible environment for its oxygen-sensitive enzymes^21^ (e.g., pyruvate synthase, 2OG synthase). Furthermore, reducing agents are available at their highest energy level in anoxic environments, and under the same environmental redox conditions, carbon fixation enzymes that use ferredoxin as a reducing agent (such as P25 in rTCA), can push this autotrophic reaction forward more than NAD(P)H types^20^, further decreasing the rTCA ATP demand. Most of the rTCA modules (leading to the production of intermediate compounds such as oxaloacetic acid (OAA), 2-oxoglutarate (2OG), and malate) were more prevalent among high GC-content lineages (**Supplementary Fig. S12**). These intermediates are key metabolites that can act as precursors for nucleotide- and amino acid-biosynthesis which will be in greater demand in high GC-content and larger genomes.

2OG and OAA interconnect carbon and nitrogen metabolism^22,23^ with 2OG being the main precursor for amino acid biosynthesis (glutamate/glutamine) and nitrogen assimilation^23^. Glutamate is generated via either the glutamine synthetase-glutamine oxoglutarate aminotransferase (GS-GOGAT) pathway (P43, P44, and P46) or the glutamate dehydrogenase (GDH) pathway (P45), both of which assimilate ammonium while using 2OG as a carbon backbone^23,24^. The GDH pathway does not require ATP, but the GS-GOGAT pathway does^25^. However, GS-GOGAT is known to be more efficient under low concentrations of 2OG or ammonium and produces two molecules of glutamate^23,24^. Both ammonium assimilation pathways were prevalent among high GC-content lineages **(Supplementary Fig. S12)**, which can potentially contribute to satisfy their higher nitrogen requirements.

At the interface of amino acids and nucleotide biosynthesis^26^, the produced glutamine can be converted to 5-phosphoribosylamine via *purF* (amido-phosphoribosyl transferase, P47) that is channeled towards purine metabolism^27^ and this enzyme also showing a higher prevalence among high GC-content lineages (**Supplementary Fig. S12**). Moreover, OAA is a precursor for the biosynthesis of aspartate and pyrimidines^22^. Modules for OAA production (P26, P27, P28), as well as those encoding its conversion to aspartate via ASPDH or aspartate aminotransferase (P29), were more prevalent in high GC-content lineages. Aspartate can also be produced from fumarate (P34) which was less prevalent than P29, yet still more common in high GC-content lineages (**Supplementary Fig. S12**).

This might imply that high GC-content MAGs/SAGs might be directing the carbon flow away from rTCA by channeling 2OG and OAA towards amino acid and nucleotide biosynthesis. However, anaplerotic pathways can compensate for such carbon flow rerouting^26^. Interestingly, some main anaplerotic pathways^28^ that can feed the rTCA cycle (e.g., carboxylation of pyruvate by pyruvate carboxylase (P28) or carboxylation of phosphoenolpyruvate by PEP carboxylase (P27); oxidation of malate to pyruvate by the malic enzyme (P32), and the glyoxylate cycle (P48, P49, 50)) were more prevalent among high GC content lineages (**Supplementary Fig. S12**). Furthermore, malate as a regulator of the central carbon metabolism^24^, can be compensated for by the glyoxylate cycle as an anaplerotic pathway. Modules within the fermentation pathways (P11, P12, P13, P20, P21) could also be connected to rTCA via their potential for producing acetyl CoA (**Fig. 2**). These modules were also more prevalent among high GC content lineages (**Supplementary Fig. S12**).

The carbon fixation step of WLP pathway (by formate dehydrogenase (P1 and P2)) or conversion of formaldehyde (P3) or n-formyl derivatives (P5) can lead to production of formate. Formate can then continue within the WLP pathway (P6, P7) and either be channeled to the reductive glycine/serine (rGly/Ser) pathway (P8, P9) or enter the pool of common goods (via passive diffusion^29^ or a formate transporter (P60) that was present in 21% of formate-producing MAGs/SAGs; **Supplementary Fig. S12**). If formate is channeled to the rGly/Ser pathway, this will cause production of glycine or serine^30^. Serine is the main C1 pool for biosynthesis of other compounds including purines and some amino acids (e.g., cysteine, methionine, and tryptophan)^31^. Alternatively, serine can also be produced via the *serB* gene (P10)^31^, which was more prevalent among high GC-content FSGD lineages (**Supplementary Fig. S12**). Serine can be converted to pyruvate via L-serine deaminase^32^, which was detected in 25.2% of MAGs with rGly/Ser pathway (32 out of 127). Alternatively, glycine can be converted to acetyl-phosphate and then pyruvate using glycine reductase complex^33^ (detected in *ca.* 8% of MAGs with rGly pathway).

FSGD MAGs/SAGs only encode the first step of 3HP pathway, producing malonyl-CoA. It is important to note that bicarbonate carboxylating enzymes in carbon fixation pathways preferentially work at substrate saturation levels and are usually active when HCO3 is available in high concentrations ^20^. This might be the reason for the lower prevalence of the 3HP pathway in FSGD MAGs/SAGs and hint at potentially low/episodic availability of HCO3 in these boreholes. However, the produced malonyl-CoA itself is a main precursor for endogenous fatty acid biosynthesis (P51) using malonyl-CoA:acyl-carrier-protein (ACP) transacylase (fabD, K00645)^34,35^. Genes involved in endogenous fatty acid biosynthesis were more prevalent among high GC-content lineages (**Supplementary Fig. S12**). FSGD MAGs/SAGs did not contain the genes needed for converting malonyl-CoA to propionyl-CoA; however, if s-methyl malonyl-CoA is present in their environment, they have the required gene to convert it to succinyl-CoA (P52) and then fumarate or 2OG (P53).

The exogenous fatty acid biosynthesis (P56), which is less energy-intensive^36^, can follow three paths via the activity of acyl-CoA synthetase (FadD), acyl-ACP synthetase (Aas), or fatty acid kinase (FakAB), to respectively produce acyl-CoA, acyl-ACP, or acyl-phosphate. These compounds are then added to glycerol-3phospate (G3P) via the activity of PlsB abd (on Acyl-CoA) and PlsXY acyltransferases (on acyl-ACP and acyl-phosphate). Then PlsC catalyzes another addition of acyl-ACP to generate phosphatidic acid, which is a precursor for phospholipidsbiosynthesis^36^. The exogenous fatty acid biosynthesis (P56) was also more prevalent among high GC-content lineages, indicating their efficient utilization of all available resources in the environment for different cellular purposes (**Supplementary Fig. S12).**

To gain a more comprehensive perspective on the distribution of metabolic strategies across the communities, the sugar-acid preference (SAP) model^37^ was used to predict the FSGD MAGs/SAGs preference. SAP ranges from 1 (extreme sugar specialists) to -1 (extreme acid specialists). The analysis showed a negative correlation between the SAP index with both GC content and EGS of FSGD MAGs/SAGs; i.e., a lower SAP index (indicative of acid preference) corresponded to higher GC content and EGS **(Supplementary Fig. S13),** potentially attributed to the presence of modules involved in metabolizing intermediates of rTCA (e.g., OAA, 2OG, malate). At module level, the density of SAP index of MAGs/SAGs for modules associated with acid consumption/production (e.g., P15, P23, P24, P31, P48, P49) and transport (specifically acetate (P79), malate (P74, P78), succinate (P74, P82), and citrate (P80, P83) import/export) were in agreement with the role of these modules in acid metabolism (**Supplementary Fig. S14**). In all boreholes, the density of the SAP index of MAGs/SAGs peaked around 0.7-0.8 (higher sugar preference); however, in OL-KR46, the peak is at around -0.5, indicating a higher acid preference of these MAGs. This further confirmed the modular metabolic analyses of this study regarding the importance of metabolizing intermediate acid compounds (i.e., organic acids such as malate, OAA, 2OG) for lineages inhabiting this borehole.

### Nitrogen acquisition-related genes were more prevalent among higher GC-content lineages

The nitrogen fixation module (PN6) was present in 78 out of 1185 FSGD genome clusters (**Supplementary Fig. S15**) and was also more prevalent among high GC-content lineages. Additionally, high-affinity nitrate/nitrite transporters (module PN1) as well as genes involved in reducing nitrate/nitrite to ammonium (modules PN2, PN3), and ammonium transporter protein (AMT, K03320)/permease (module PN7) were more prevalent among high-GC lineages. Higher GC content means higher nitrogen requirements since guanine and cytosine pairs require one more nitrogen than adenine and thymidine pairs, which could potentially explain the higher prevalence of different nitrogen uptake and fixation strategies in lineages with higher GC content (**Supplementary Fig. S15**).

### Shared metabolic networks in different boreholes offer similar niches that are filled by different lineages

Communities of different boreholes represented a similar distribution profile for different metabolic modules (**Fig. 3**). This imply that similar metabolic niches were available in the different boreholes, but that lineages with metabolic capabilities that can occupy these niches and the range of their genome GC-content varied across the different boreholes (**Fig. 4 & Supplementary Fig. S16**).

**Figure 3:**
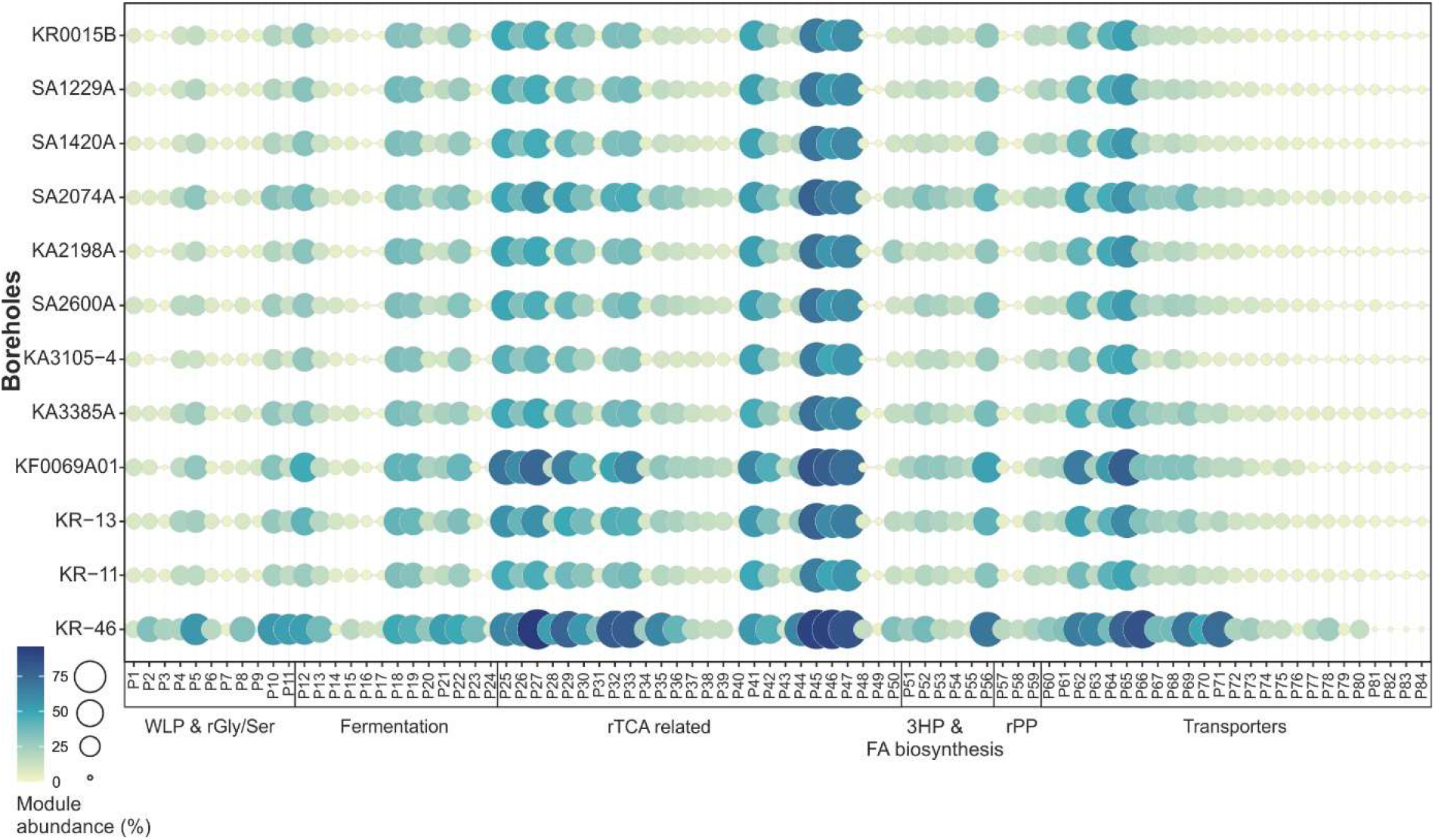
Prevalence of metabolic modules in different boreholes. The size and color of circles indicate the percentage of members within each borehole possessing a corresponding metabolic module. Boreholes are arranged based on their depth from the top to the bottom of the plot.

**Figure 4:**
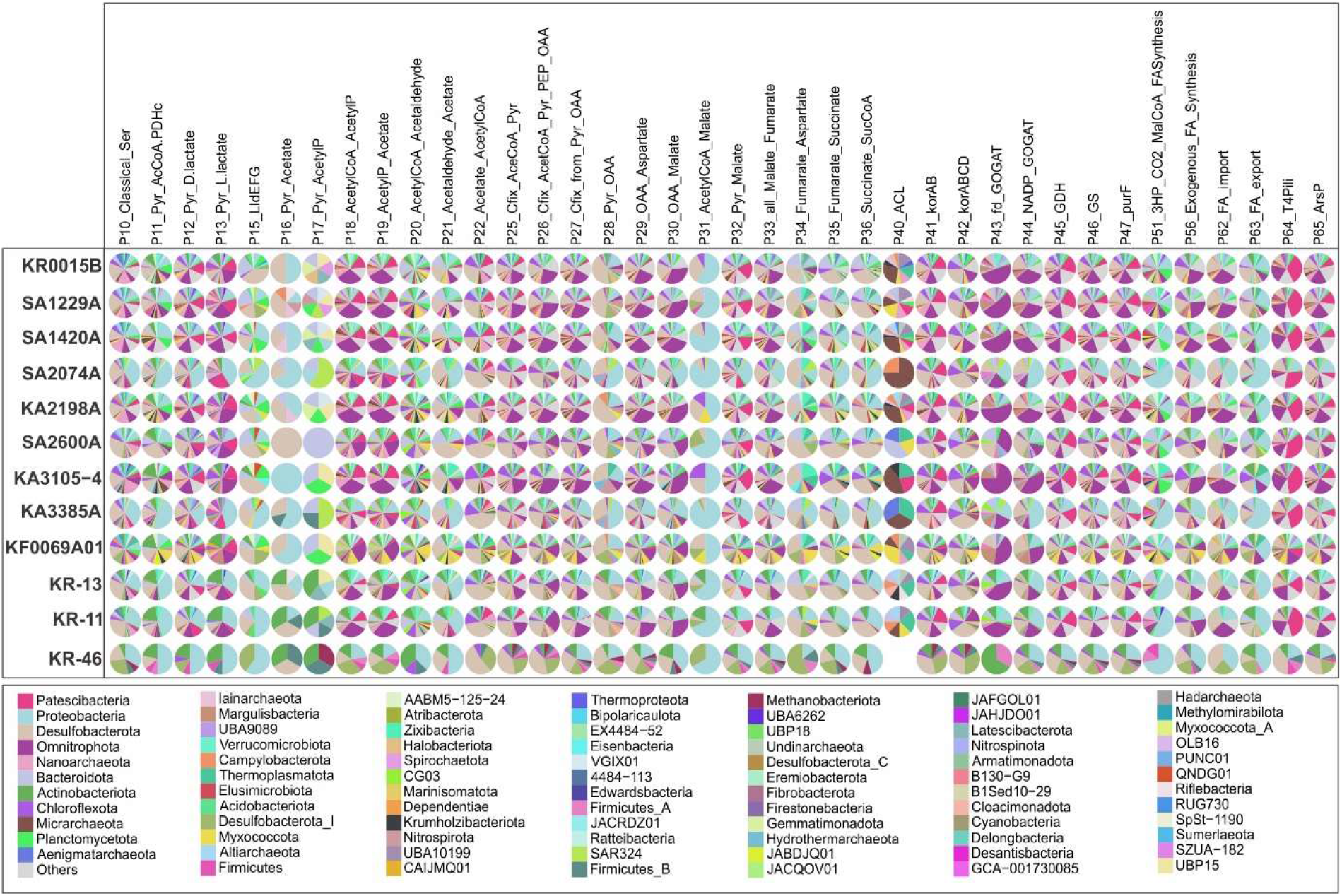
Composition of microbes carrying genes for each core metabolic module in different boreholes. The community of microbes encoding each metabolic module in different boreholes are shown as a pie chart. The color legend at the bottom of the plot shows taxonomic affiliation of MAGs/SAGs encoding different modules at phylum level. Phyla with overall abundance lower than 1% in all boreholes were clumped together as “others” category.

For the entire TCA cycle to run in the carbon fixation direction (rTCA) three enzymes, named fumarate reductase (P35), 2-oxoglutarate: ferredoxin oxidoreductase (2OGOR or *korABCD* respectively modules P37 and P42), and ATP citrate lyase (ACL, module P40), must be present^38^ (**Fig. 5**). Taking this into account, only eight FSGD MAGs/SAGs (affiliated to Campylobacterota, Bacteroidota, Myxococcota, Desulfobacterota and Thermoplasmatota phyla) encoded all the genes to carry out rTCA. However, none of these lineages were detected in boreholes OL-KR46 and KA3385A **(Supplementary Table S4)**.

**Figure 5:**
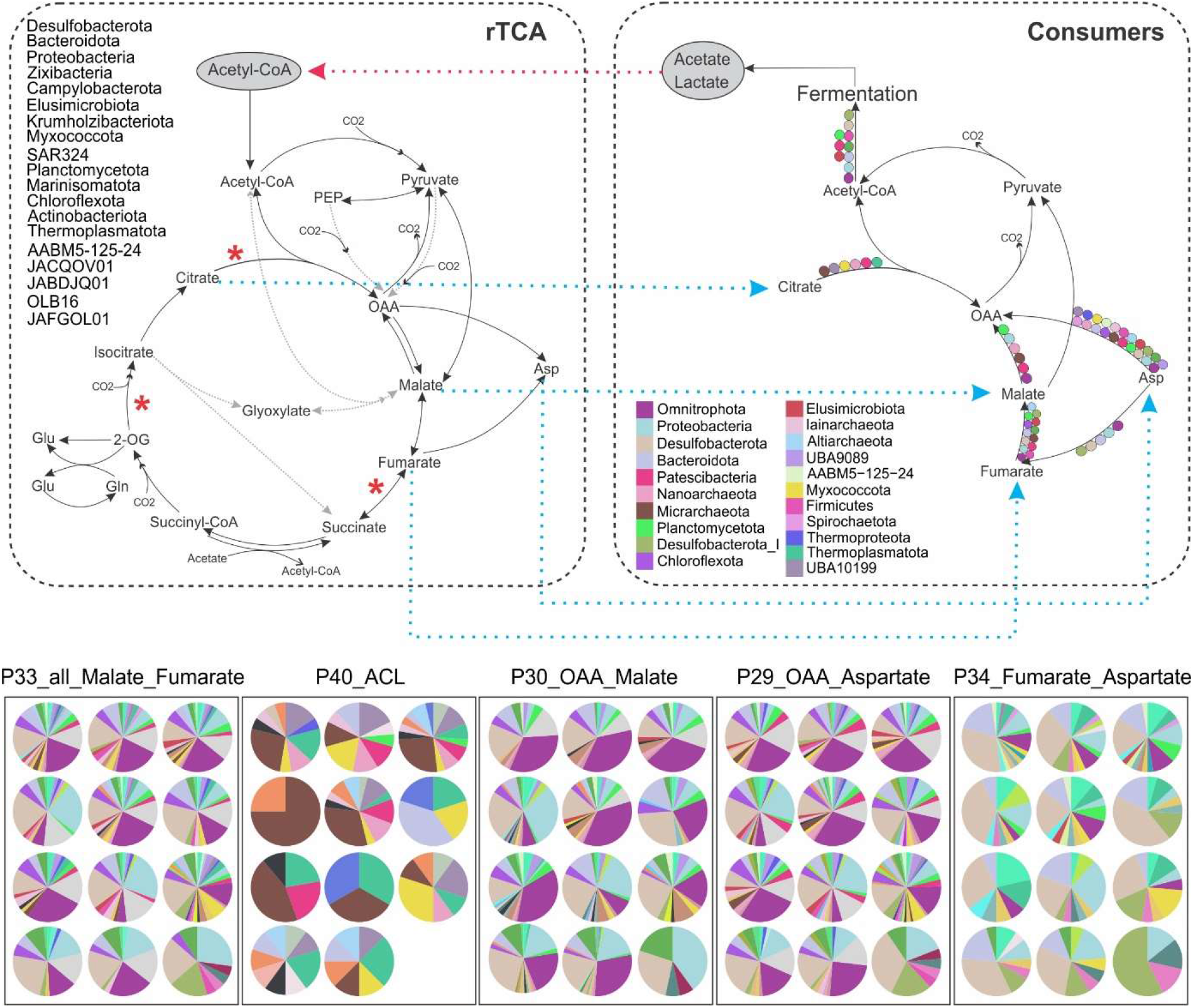
Metabolic cross-feeding among microbial communities of different boreholes. Top left panel shows the complete rTCA cycle, highlighting phyla encoding at least two of the three genes critical for rTCA to operate in the carbon fixation direction (denoted by an asterisk). Top right panel showcases the truncated TCA pathway detected in consumers. The main phyla encoding each module are represented in circles with the color legend included in the bottom left side of the panel. Blue dashed arrows show the potential transfer of metabolic products from primary producers to consumers. The red dashed arrow indicates the potential reciprocal utilization of products by primary producers. Bottom panel denotes the composition of microbes carrying each of the metabolic modules present in the consumers in this cross feeding in different boreholes as pie charts. In each subpanel, boreholes are arranged from left to right based on their depth. In bottom subpanels, the order of boreholes from right to left is based on depth as follows; Row 1: KR0015B, SA1420A, SA2074A; Row 2: KA2198A, SA1229A, KF0069A01; Row 3: SA2600A, KA3385A, KA3105-4; Row 4: KR-11, KR-13, KR-46

There were 113 additional MAGs that carried two out of the three necessary enzymes (P35, P37). This could be either due to MAG incompleteness or that they are only performing carbon fixation up to the production of citrate. These lineages were affiliated with Desulfobacterota, Proteobacteria, Bacteroidota, Campylobacterota, Zixibacteria, Elusimicrobiota, Myxococcota, Krumholzibacteriota, Planctomycetota, SAR324, Thermoplasmatota, AABM5-125-24, Actinobacteriota, Chloroflexota, Marinisomatota, JABDJQ01, JACQOV01, OLB16, and JAFGOL01 phyla (**Figs. 4&**5**, Supplementary Fig. S17 & Table S4**). From these MAGs, four were present in the OL-KR46 borehole (affiliated to Desulfobacterota and Actinobacteriota). The GC content of lineages carrying rTCA-specific genes ranged from 30.88 to 68.26, and their EGS ranged from 2 to 10 Mb (**Supplementary Table S4**). However, from across the range of GC content and EGS, these primary-producer lineages can develop large population sizes (**Supplementary Fig. S17**).

Many of the studied modules lead to the production of intermediate compounds such as malonyl-CoA, OAA, 2OG, and fumarate. These intermediates can either be channeled to different cellular pathways (e.g., biosynthesis of fatty acids, amino acids, and nucleotides) or be released to the environment either by diffusion through the cell membrane or via specialized transporters (for compounds overproduced in the cell or those that cannot be further metabolized). If these compounds were to be released into the environment, they become a part of a public goods pool, potentially contributing to promiscuous metabolic cross-feeding in the deep oligotrophic groundwater^39^.

Even if MAGs/SAGs carry only few enzymes relevant for metabolizing some of these intermediates (rather than featuring the complete pathway), they can contribute to the flow of carbon within and between cells. For example, MAGs/SAGs carrying modules P25 to P33 (i.e., a truncated TCA cycle) in each borehole could take up and metabolize fumarate (as an example of a public good) and even transform it to acetyl-CoA (in reverse from module P33 towards P25). These MAGs/SAGs (*n*=367) were mainly affiliated to Omnitrophota, Proteobacteria, Desulfobacterota, Bacteroidota, Chloroflexota, Planctomycetota, Actinobacteriota, Micrarchaeota, Patescibacteria, and Nanoarchaeota phyla (**Figs. 4&**5). A number of these MAGs/SAGs (*n*=190) also carried genes for PAT-ACK (P18 and P19; specifically *n*=64 belonged to phylum Omnitrophota), which further converts acetyl-CoA into acetate while also generating one ATP. The produced acetate could be released to the surrounding environment via the formate transporter (P60)^40^ that was present in 51 of these 190 fermenter lineages (**Supplementary Fig. S16**), with 22 of them belonging to the phylum Omnitrophota. Acetate, in turn, can be taken up by primary producers and transformed into acetyl-CoA (via P22 or P23) to replenish the rTCA pathway in an intricate and promiscuous community-level cross-feeding. Acetate could also be turned into pyruvate (P16), which feeds the rTCA (**Fig. 4**).

Representatives from Patescibacteria (Portnoybacterales, BM507, SG8-24, JAHISY01, UBA1369), Nanoarchaeota (Pacearchaeales, Woesearchaeales), and Micrarchaeota (Norongarragalinales, Anstonellales, JACRGF01) can further use promiscuous transporters (e.g., P64, P65) to take up malate from the pool of “public goods” (**Fig. 4**) and convert it to pyruvate via malate dehydrogenase (P32) while also conserving reducing power by producing NADPH^41^. 21.7 % of the lineages encoding enzymes for P64/P65 are affiliated to Patescibacteria. Lineages carrying P32 can subsequently turn the produced pyruvate into fermentation products such as D-lactate/L-lactate (P12, P13; **Fig. 4**) and release it into the environment. Interestingly, 34 of the primary producer lineages encoding rTCA utilized lactate via lactate dehydrogenase (LldEFG^42^, P15; **Fig. 4**). The activity of this lactate dehydrogenase leads to the production of pyruvate and feeds in to the rTCA cycle. This community-level network of reciprocal metabolic cross-feeding was detected in all boreholes, although at different prevalence and in different microbial lineages (**Figs. 4&5 and Supplementary Fig. S16)**.

Apart from this main network of metabolic cross-feeding around intermediates of the rTCA pathway, some other rTCA pathway intermediates can shape peripheral cross-feeding nodes. For example, most MAGs/SAGs with rTCA did not contain ACL (P40) to regenerate the substrates of the cycle. However, they can release citrate to the surrounding environment, which can be taken up by ACL-bearing MAGs and be converted to acetyl-CoA and OAA. This potential was mainly present in representatives of Micarchaeota, UBA10199, Nanoarchaeota, Thermoplasmatota, Myxococcota, Patescibacteria, Undinarchaeota and Altiarchaeota (**Figs. 4&**5). However, ACL was not detected in any of the OL-KR46 MAGs.

Some MAGs/SAGs encoded the capacity to take up aspartate from the pool of common goods and convert it to malate/fumarate (P29/P34; *n*=422), which is then channeled towards pyruvate production. These lineages mainly belonged to Omnitrophota (Koll11), Proteobacteria, Desulfobacterota, Planctomycetota, Patescibacteria, Bacteroidota, Chloroflexota, Micrarchaeota, and Nanoarchaeota phyla. Moreover, these MAGs/SAGs can carry out fermentation (P18 and P19) and provide acetate to the community (*n*=214; **Figs. 4&**5). Overall, bidirectional interactions are shown to be more prevalent under anoxic conditions where they enable the community to survive in nutrient- poor environments^39^.

Finally, the 3HP pathway provides intermediates that can be directed toward FA biosynthesis (P51). Genes encoding this module were present among representatives of Proteobacteria, Bacteroidota, Planctomycetota, Zixibacteria, Firmicutes, AABM5-125-24, Omnitrophota (Koll11), and Elusimicrobiota (**Fig. 4**). Moreover, 62 of the 170 lineages encoding this module also encoded genes for fatty acid export (using AcrAB-TolC complex or FarE^43^, P63; **Supplementary Fig. S16**) and were detected in all boreholes. These lineages can potentially supply fatty acids to CPR representatives that are not capable of synthesizing fatty acids. Surveyed FSGD lineages did not encode the full 3HP pathway and only contain acetyl-CoA carboxylase that produces malonyl-CoA which supplies the fatty acid biosynthesis.

## Conclusion

We observed decreased metabolic cross-feeding potential due to low cell density affect the eco-evolutionary trajectory of microbes in deeper groundwater. The prevalence of lineages such as DPANN and Patescibacteria with patchy metabolism that are most often reliant on symbiotic relations, decreased in deeper and more oligotrophic groundwaters. Furthermore, lineages with larger genome sizes (and higher GC content) appear to build up larger populations in such ecosystems. Detailed analyses of metabolic modules in different boreholes showed that similar niches were available not only for primary producers but also for coupled cross-feeding networks. However, different populations filled these shared niches in different boreholes. The study highlights the decisive role of metabolic cross-feeding for genome evolution and community assembly in deep oligotrophic groundwaters and advances our understanding of the complex interactions and evolutionary pressures in these largely hidden ecosystems on our planet. The results further emphasize that mainstream theories need to be carefully vetted before being applied to new ecosystems with potentially different eco-evolutionary constraints.

## Methods

### Fennoscandian Shield Genomic Database

An extensive “multi-omics” dataset of carbon and energy-limited deep groundwater samples collected from the Swedish Nuclear Fuel and Waste Management Company (SKB) managed Äspö Hard Rock Laboratory (Äspö HRL) in Sweden and drillholes in Olkiluoto Island, Finland operated by Posiva Oy was used in this study. This dataset contained 43 metagenomes and 114 SAGs (**Supplementary Table S3**)^3^. Metagenomes originate from samples collected from nine boreholes in Äspö HRL (KR0015B, SA1229A, SA1420A, SA2074A, KA2198A, SA2600A, KA3105-4, KA3385A, KF0069A01) and three drillholes in Olkiluoto Island (OL-KR13, OL-KR11, OL-KR46). Samples were collected from different depths in the range of 70 to 528 mbsl (details and metadata for all metagenomes are shown in **Supplementary Table S1**).

Metagenome assembled genomes (MAGs) were reconstructed following the method described by Mehrshad et al.^3^. Briefly, the metagenomic sequences were quality-checked and trimmed using Trimmomatic (version 0.36)^44^. The Illumina TruSeq adapter was trimmed based on specific parameters (’TruSeq3-PE-2.fa:2:30:15 LEADING:3 TRAILING:3 SLIDINGWINDOW:4:15 MINLEN:31’). Each dataset was then individually assembled using MEGAHIT (version 1.1)^45^ with customized settings (--k-min 21 - k-max 141 --k-step 12 --min-count 2). Following assembly, contigs of at least 2 kb in length were automatically binned using MetaBat2^46^ with its default settings.

Genome-resolved analyses of these metagenomes and single cell amplified genomes resulted in the reconstruction of a total of 1990 MAGs/SAGs with ≥50% completeness and ≤5% contamination according to CheckM (v.1.2.0)^47^ assessment. This database of MAGs/SAGs reconstructed from these metagenomes and used in this study is referred to as the “Fennoscandian Shield Genomic Database” (FSGD for short). The taxonomic affiliation of reconstructed MAGs/SAGs was assigned using GTDB-tk v2.1.0 (reference database R207)^48^ .

### MAGs/SAGs abundance calculation in different metagenomes

All MAGs and SAGs were further clustered into mOTUs (metagenomic operational taxonomic units) using mOTUlizer (v.0.3.2) at the 95% average nucleotide identity (ANI) threshold^49^. Representative MAGs or SAGs of each mOTU were used for abundance calculation. The abundance of representative MAGs/SAGs in each metagenome was calculated with CoverM (v.0.6.1) using TPM as the normalization method (https://github.com/wwood/CoverM).

### Functional annotation of reconstructed MAGs/SAGs and modular metabolic analyses

FSGD MAGs and SAGs were annotated using PROKKA (v.1.12)^50^, and then functions were assigned using eggNOG-mapper (v.2.0.15)^51^. In the next step, the presence/absence of genes involved in six prevalent carbon fixation pathways (i.e., reductive citrate cycle (rTCA), reductive acetyl-CoA/Wood-Ljungdahl pathway (WLP), phosphate acetyltransferase-acetate kinase (PAT-ACK), Calvin-Benson-Bassham cycle/reductive pentose phosphate cycle (rPP, rPP-Calvin), 3-hydroxypropionate bi-cycle (3HP) and dicarboxylate-hydroxybutyrate cycle) in different MAGs/SAGs was surveyed using a modular approach as explained below. Additionally, the presence/absence of genes involved in nitrogen metabolism were also surveyed.

To gain a better insight into how these pathways affect the microbial community and their metabolic interdependencies, the C-fixation and nitrogen acquisition pathways were subdivided into modules that lead to the production of intermediate compounds (e.g., formate, pyruvate, oxaloacetate, etc.; **Fig. 2**). The list of KEGG orthologs (KOs) for genes involved in each module is provided in **Supplementary Table S3**. In total, 58 carbon fixation-related modules and 25 transporter genes were inspected. Similarly, modules involved in nitrogen acquisition and assimilation and their related transporters were investigated (**Supplementary Table S3**).

The prevalence of these modules/genes was analyzed in the range of genome size and genome GC content. In order to normalize the data for comparative analyses, the GC content range (25-75%) of FSGD MAGs/SAGs was divided into five intervals and the prevalence of each module/gene was calculated by dividing the number of MAGs containing the module/gene in each interval by the total number of MAGs/SAGs in that interval. The same approach was applied for genome size by dividing the range of genome size (0.6 to 10.3 mb) into ten equal intervals.

### Sugar-Acid Preference (SAP) analyses

To predict the SAP for MAGs, the procedure outlined in Ref. 36^37^ was followed. Briefly, the total abundance (normalized by number of genes) of a list (given in the SI of Ref. 36) of sugar (*S*) and acid (*A*) genes was computed. The relative abundances of S and A were then used to predict SAP as tanh(s*S* + a*A*), with s = 60.76 and a =−20.21.

## Data availability

Previously released FSGD MAGs can be accessed through the NCBI BioProject under the accession number PRJNA627556. SAGs are publicly available in figshare with the identifier https://doi.org/10.6084/m9.figshare.12170313 under the project name ‘Fennoscandian Shield genomic database (FSGD)’. The additional MAGs generated for this study are deposited under the BioProject accession number PRJNA1023754 and their accession numbers are given in **Supplementary Table S1**.

## Author contributions

MD, MM, and SB devised the study. MRS and MM performed the bioinformatics analysis with contributions from CG and MG. ST was involved in the sampling of new datasets. MRS and MM interpreted the results and drafted the manuscript with input from MD. All authors read, commented on, and approved the final manuscript.

## Acknowledgments

The Swedish Nuclear Fuel and Waste Management Co (SKB) is acknowledged for providing access to the Äspö HRL and Sicada database. Maryam Rezaei Somee was supported by a scholarship from the Wenner Gren Foundation (grant no. UPD2021-0082) awarded to Mark Dopson. Maliheh Mehrshad was supported by a grant from the Swedish Research Council for Sustainable Development; FORMAS (grant no. 2021-00546). Mark Dopson was supported by The Swedish Research Council (Vetenskapsrådet contract 2018-04311). Carolina González-Rosales was supported by The Olle Engkvist Foundation (contract 216-0462) awarded to Mark Dopson. Bioinformatics analyses were carried out utilizing the Uppsala Multidisciplinary Center for Advanced Computational

Science (UPPMAX) at Uppsala University (projects NAISS 2023/22-893, 2023/6-261, and 2024/5-56). The computations were enabled by resources provided by the Swedish National Infrastructure for Computing (SNIC) at UPPMAX partially funded by the Swedish Research Council through grant agreement no. 2016-07213. Funding for open-access publishing is provided by Linnaeus University.

---

**Supplementary Figure S1:**
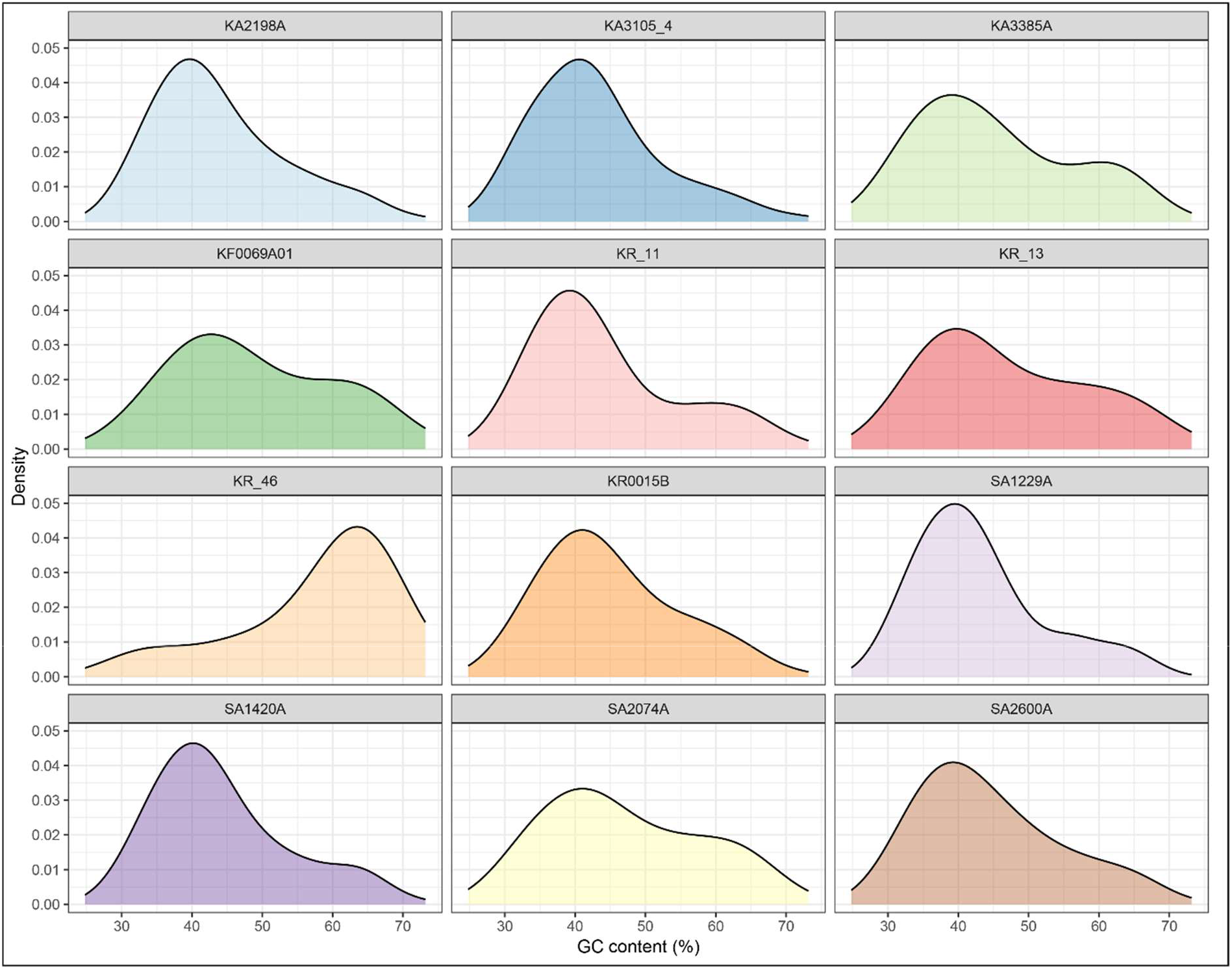
GC density of representative MAGs/SAGs present in different boreholes. MAGs/SAGs with nonzero log10 value of the calculated transcript per million (TPM) were considered as present.

**Supplementary Figure S2:**
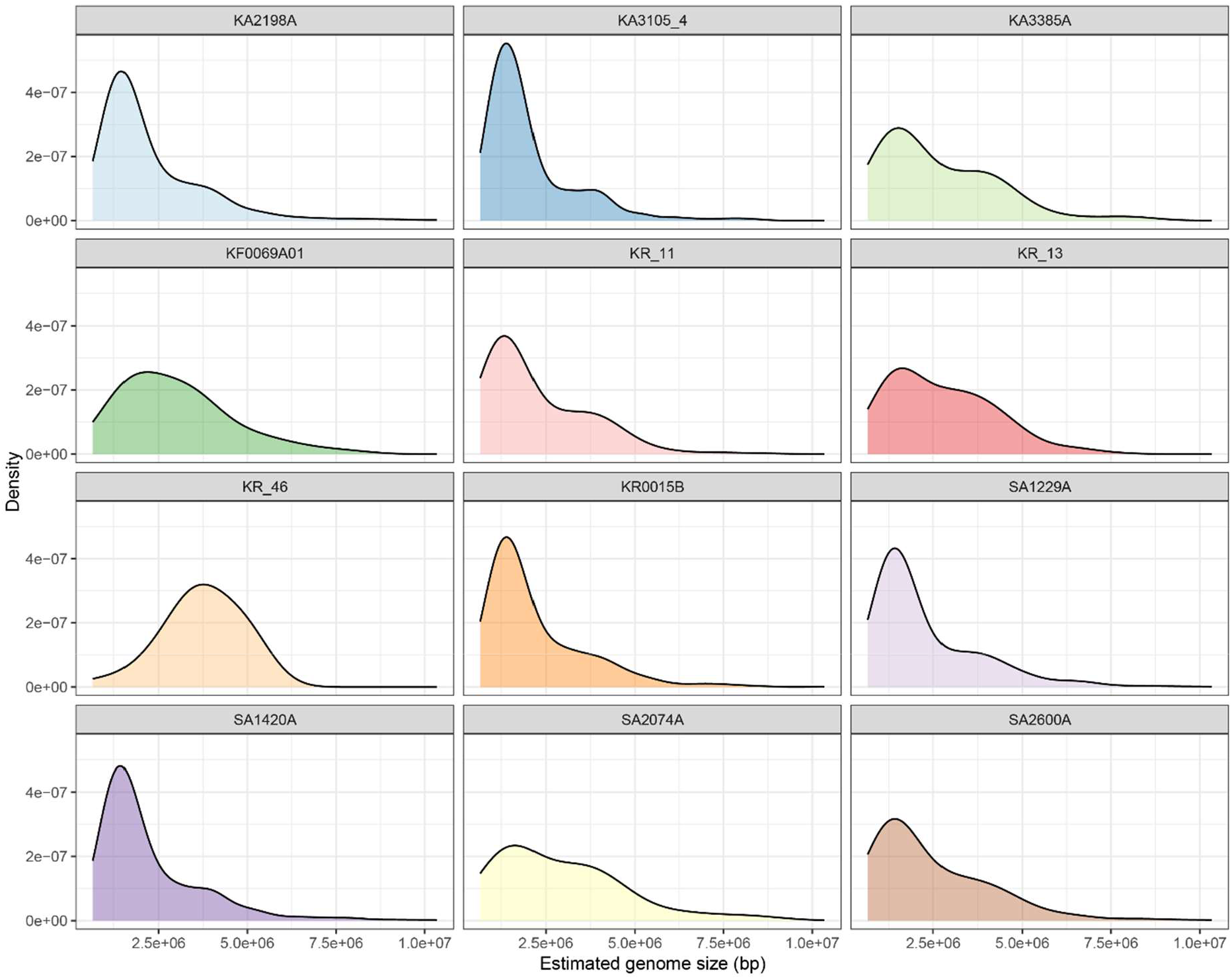
Estimated genome size (EGS) density of representative MAGs/SAGs present in different boreholes. MAGs/SAGs with nonzero log10 value of the calculated transcript per million (TPM) were considered as present.

**Supplementary Figure S3:**
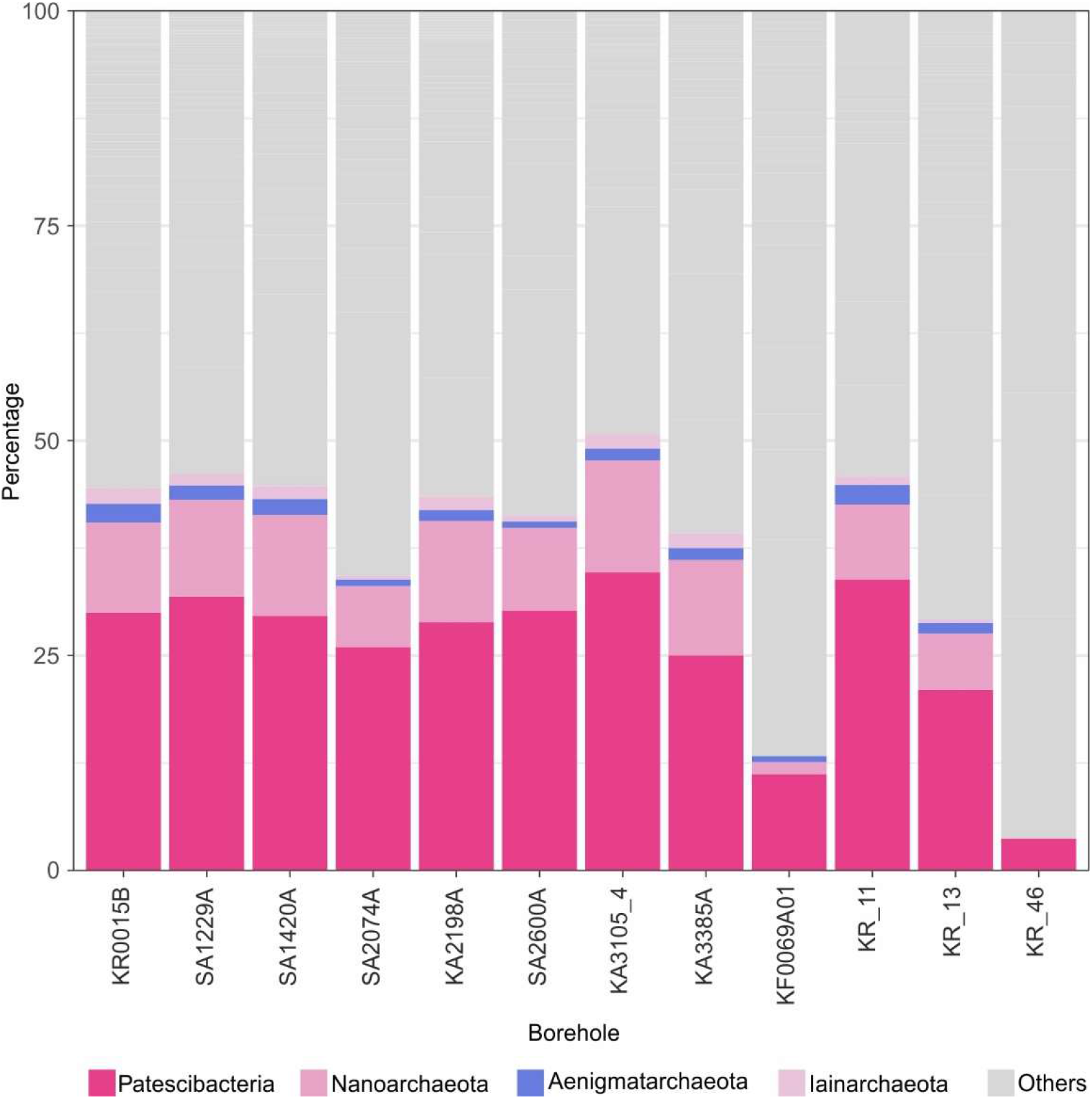
The percentage of MAGs/SAGs affiliated to DPANN superphylum and Patescibacteria phylum among MAGs/SAGs present in different boreholes.

**Supplementary Figure S4:**
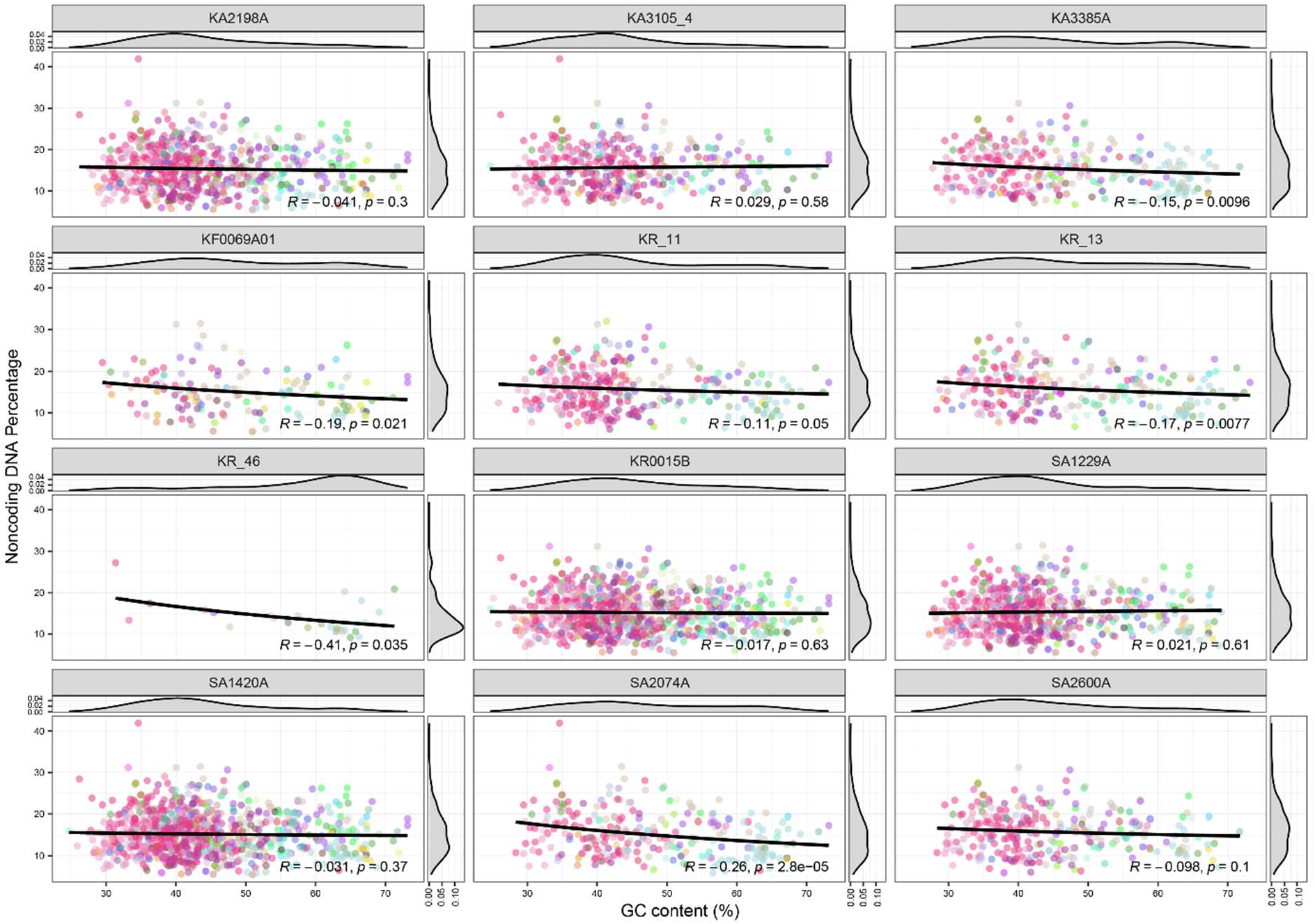
Percentage of non-coding DNA in representative MAGs/SAGs in correlation with their GC content.

**Supplementary Figure S5:**
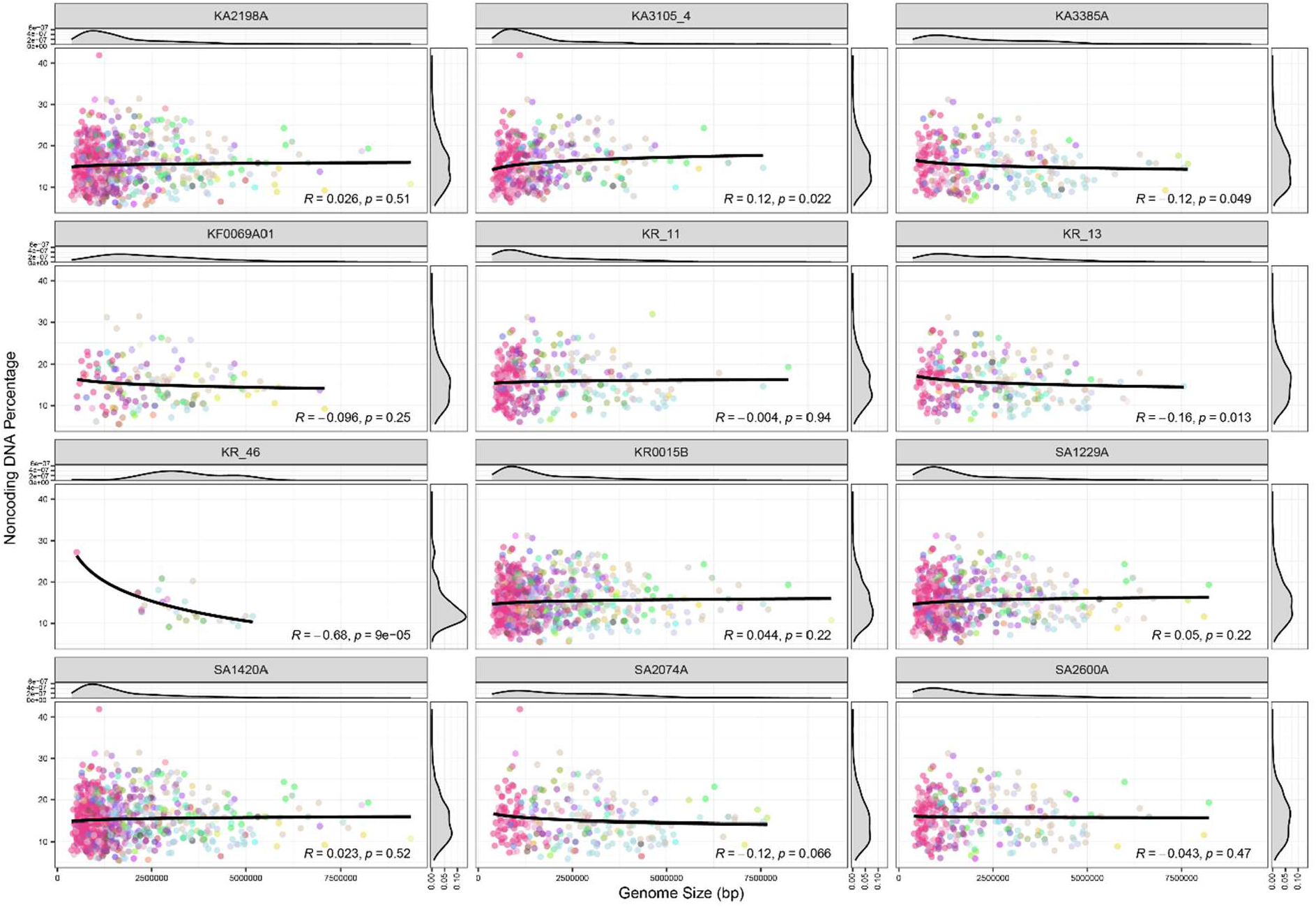
Percentage of non-coding DNA in representative MAGs/SAGs in correlation with their genome size.

**Supplementary Figure S6:**
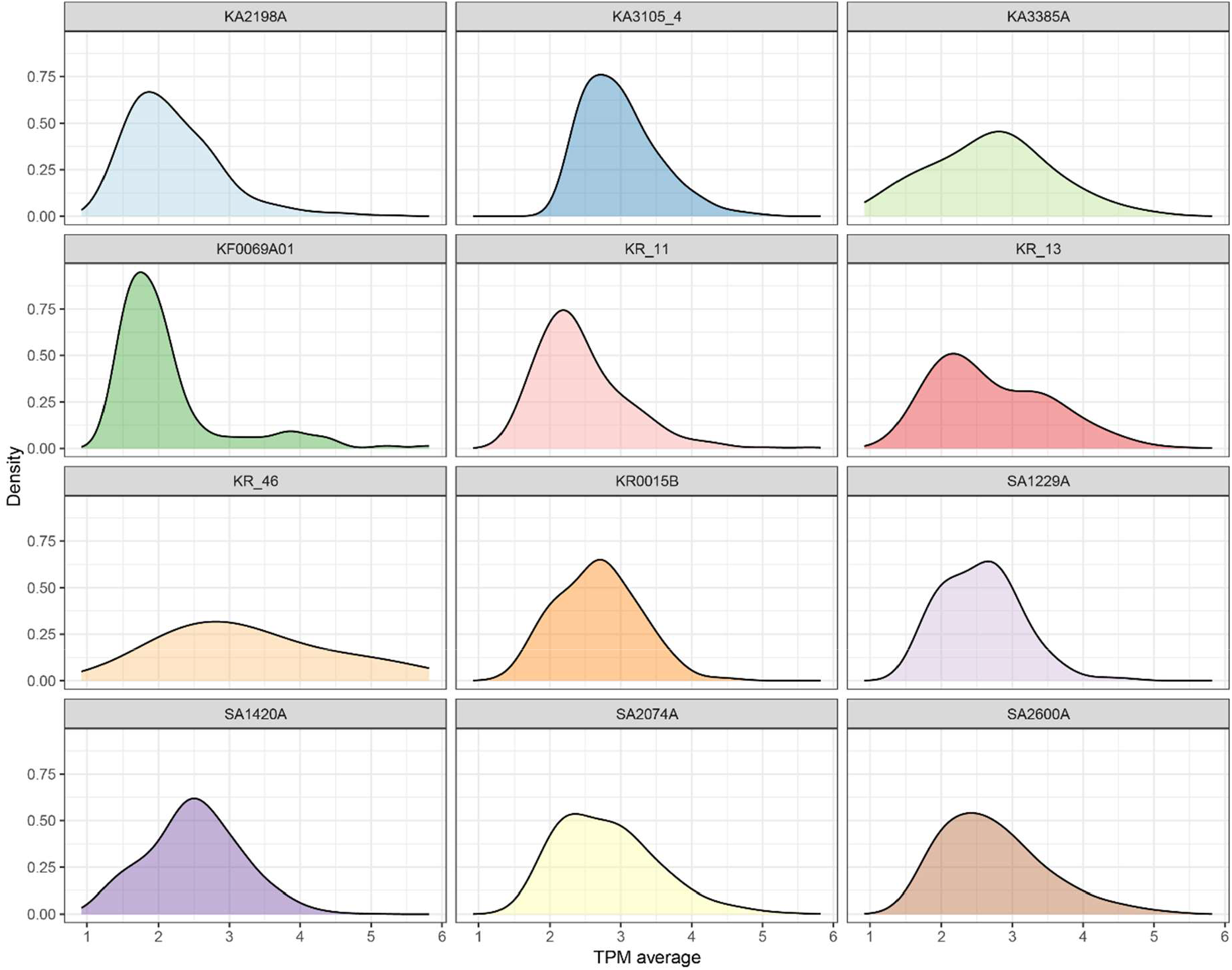
Population size distribution of MAGs in different boreholes. Population size was calculated as the average of the nonzero log10 TPM value of each representative MAG in all metagenomes sequenced for each borehole.

**Supplementary Figure S7:**
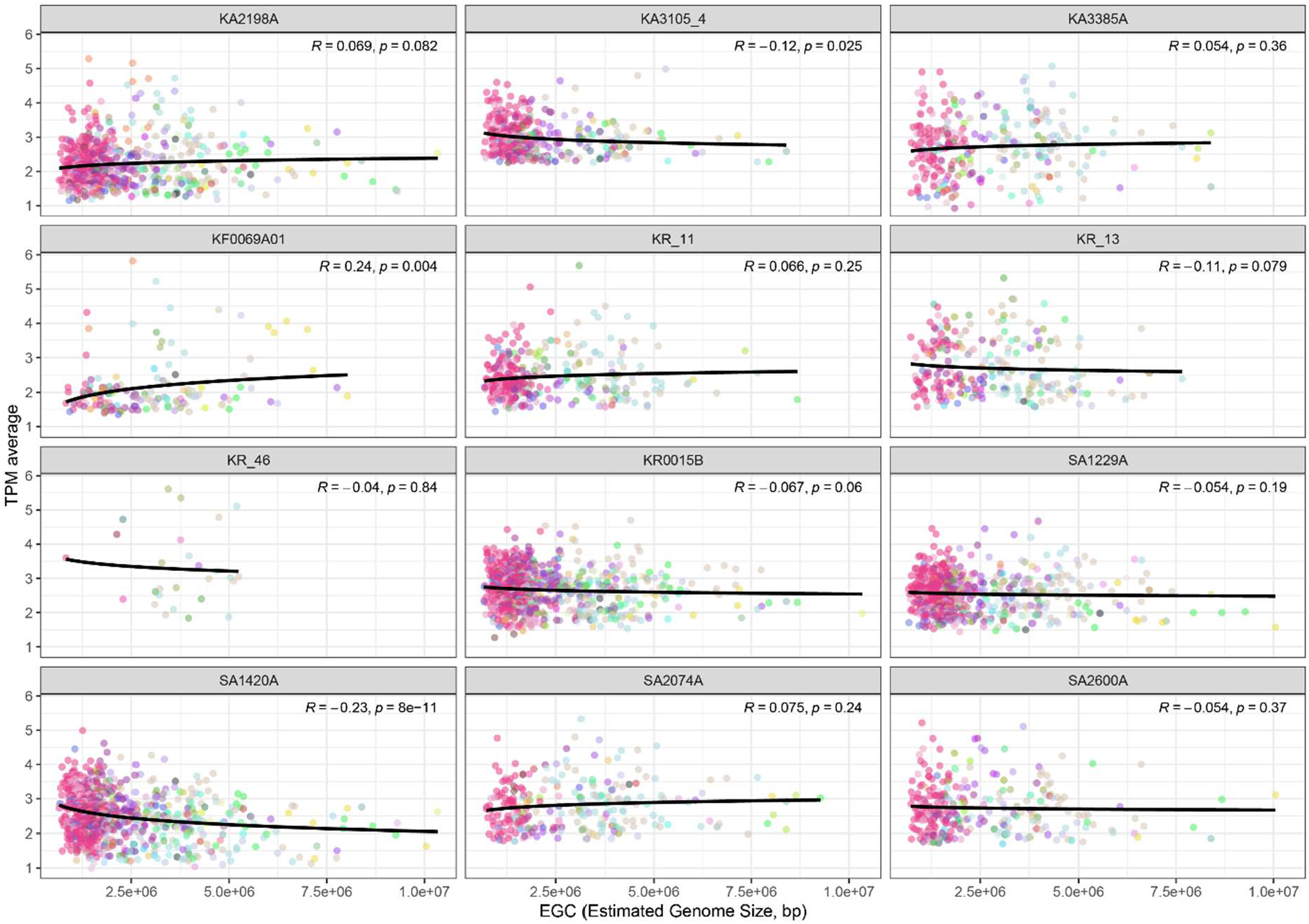
Average abundance of representative MAGs/SAGs present in different boreholes across the range of estimated genome size (EGS).

**Supplementary Figure S8:**
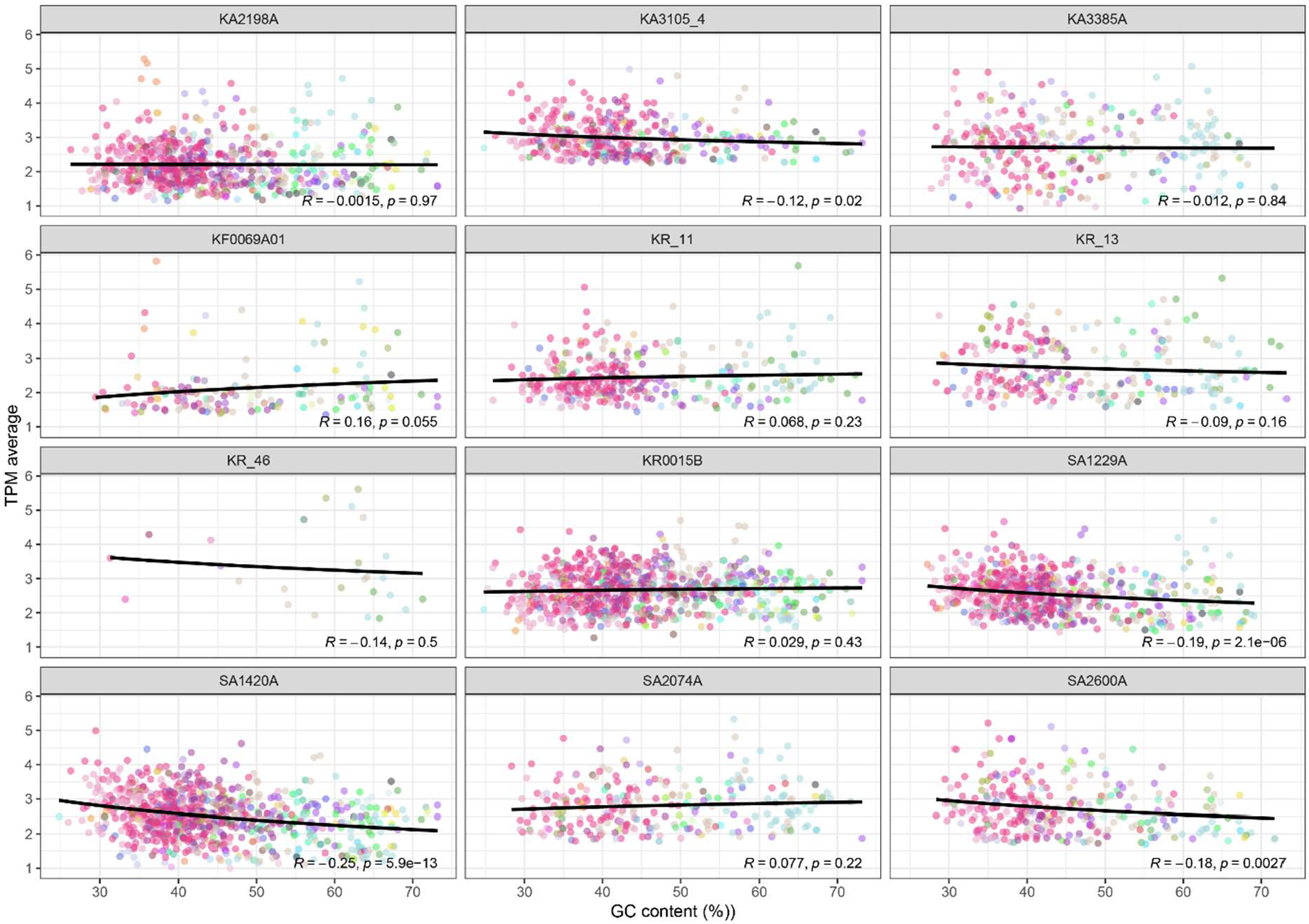
Average abundance of representative MAGs/SAGs present in different boreholes across the range of GC content.

**Supplementary Figure S9:**
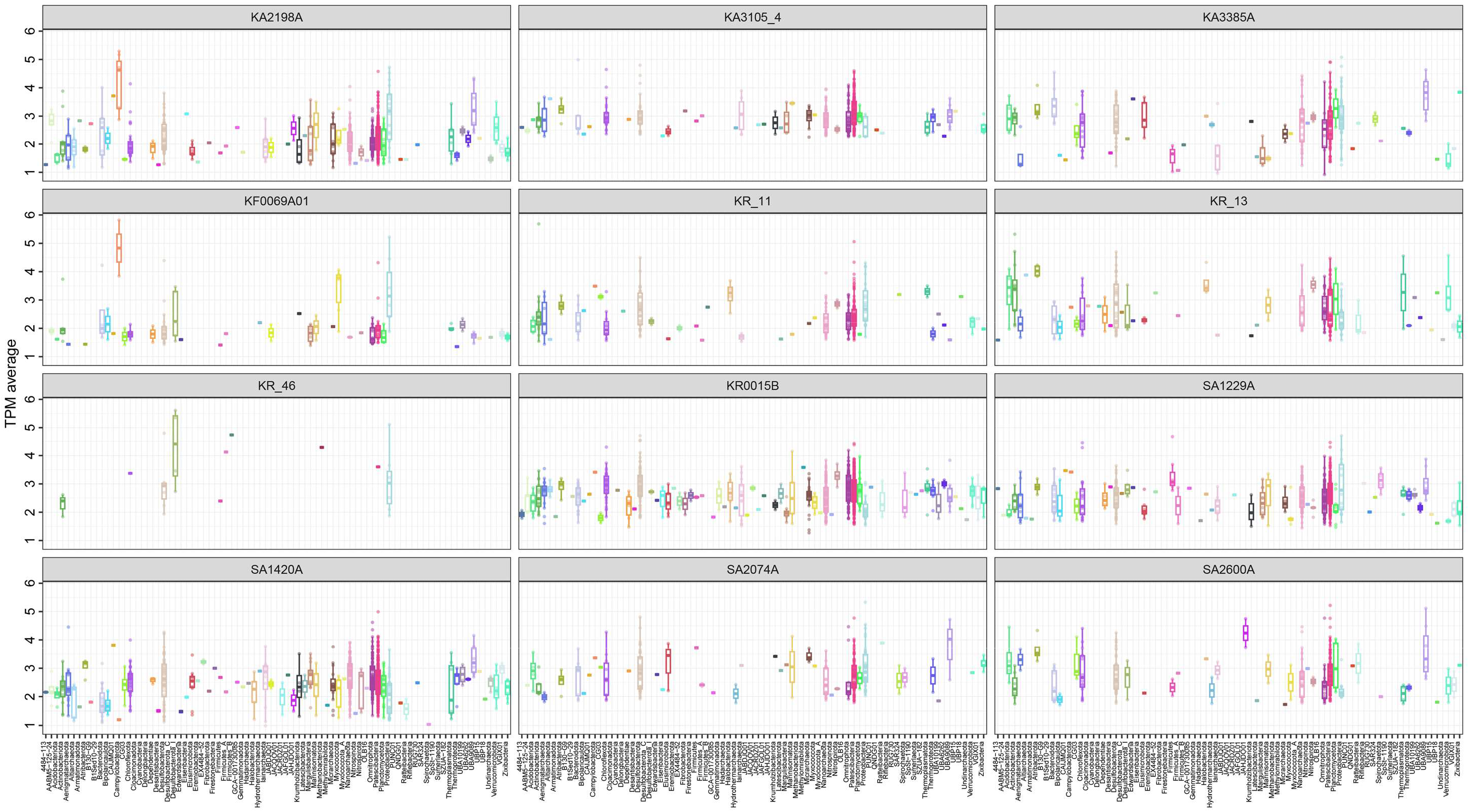
The distribution of population size for different phyla present in the boreholes. Population size was calculated as the average of the nonzero log10 TPM value of each representative MAG in all metagenomes sequenced for each borehole.

**Supplementary Figure S10:**
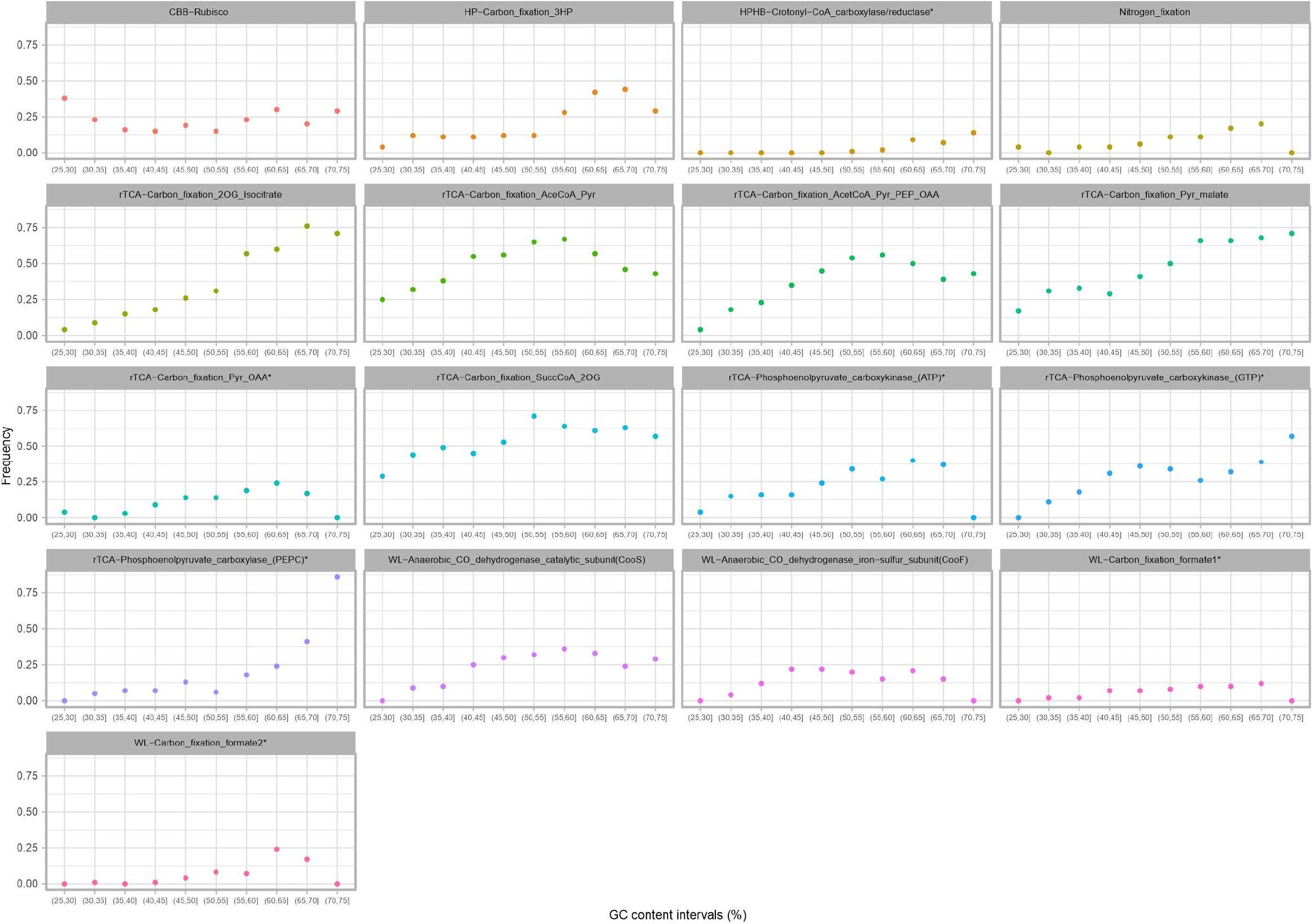
The frequency of MAGs/SAGs containing genes for different carbon fixation pathways and nitrogen fixation pathway across the range of GC content. The number of MAGs/SAGs containing genes for each pathway was counted in intervals of 5% GC content and then normalized by the number of all MAGs/SAGs present in each interval.

**Supplementary Figure S11:**
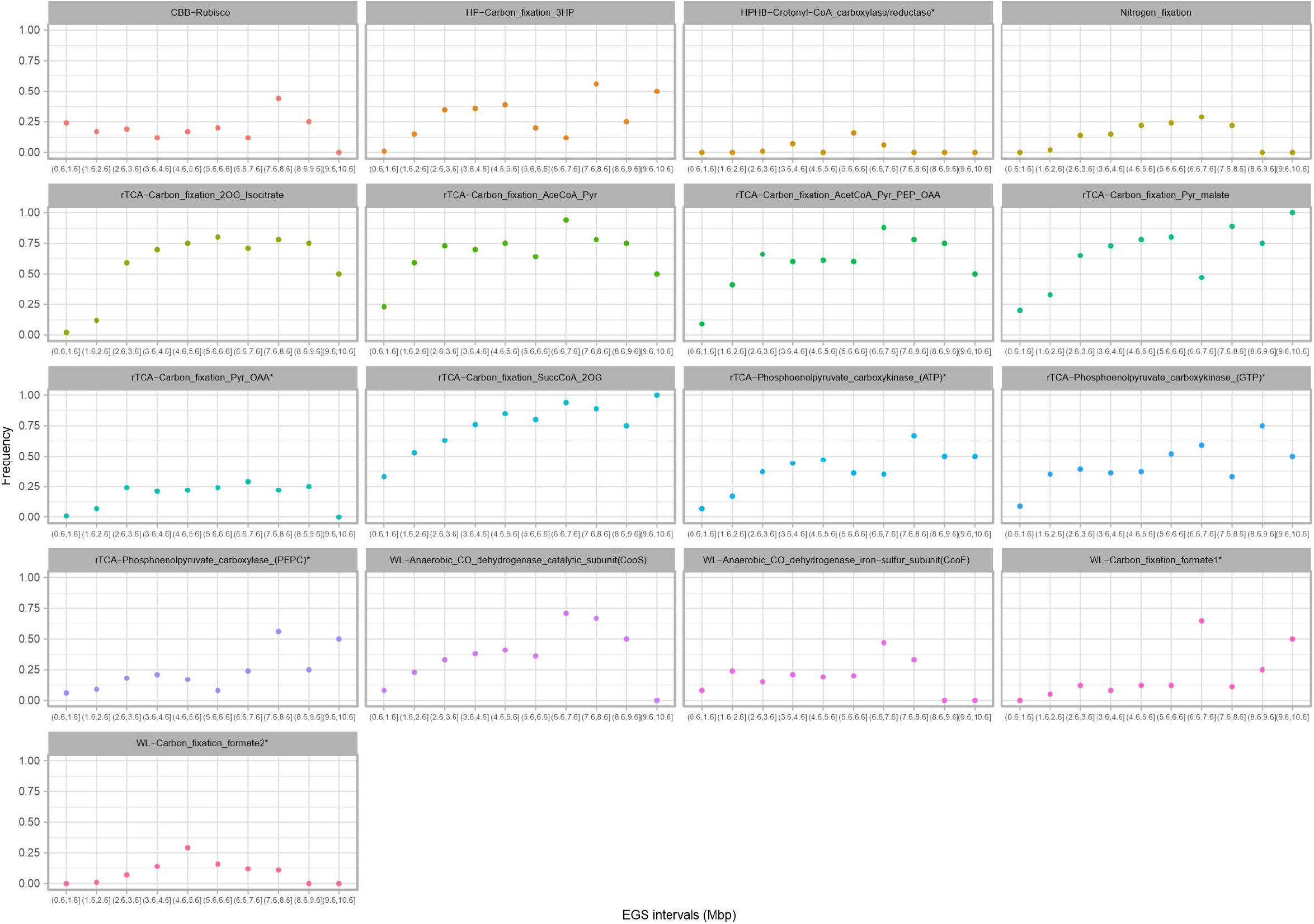
The frequency of MAGs/SAGs containing genes for different carbon fixation pathways and nitrogen fixation pathway across the range of estimated genome size. The number of MAGs/SAGs containing genes for each pathway was counted in intervals of 1 Mbp genome size and normalized by the number of all MAGs/SAGs present in each interval.

**Supplementary Figure S12:**
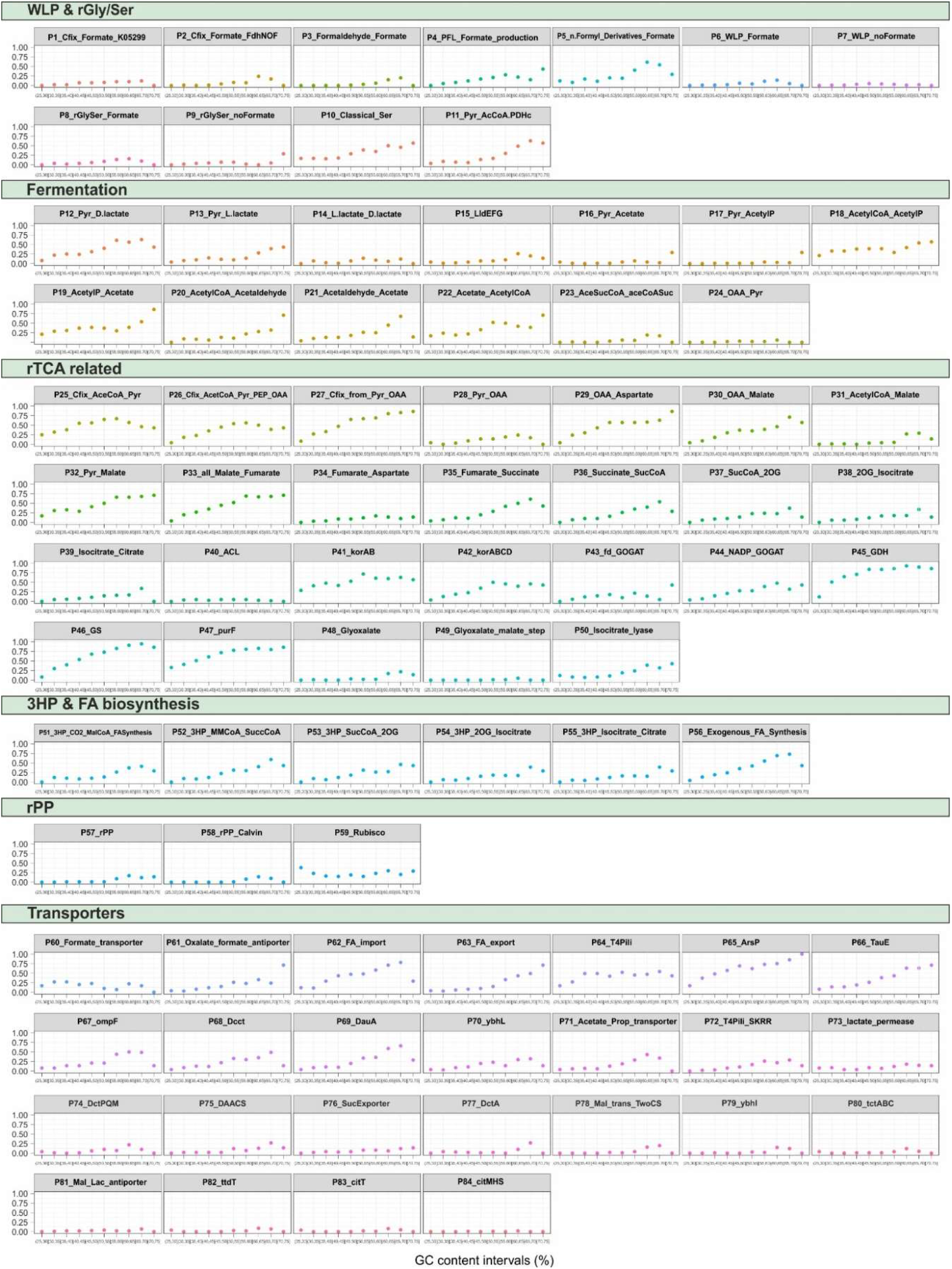
The frequency of MAGs/SAGs containing genes for 58 modules in carbon fixation metabolism as well as genes encoding 25 transporters across the range of GC content. The number of MAGs/SAGs containing genes for each module was counted in intervals of 5% GC content and normalized by the number of all MAGs/SAGs present in each interval.

**Supplementary Figure S13:**
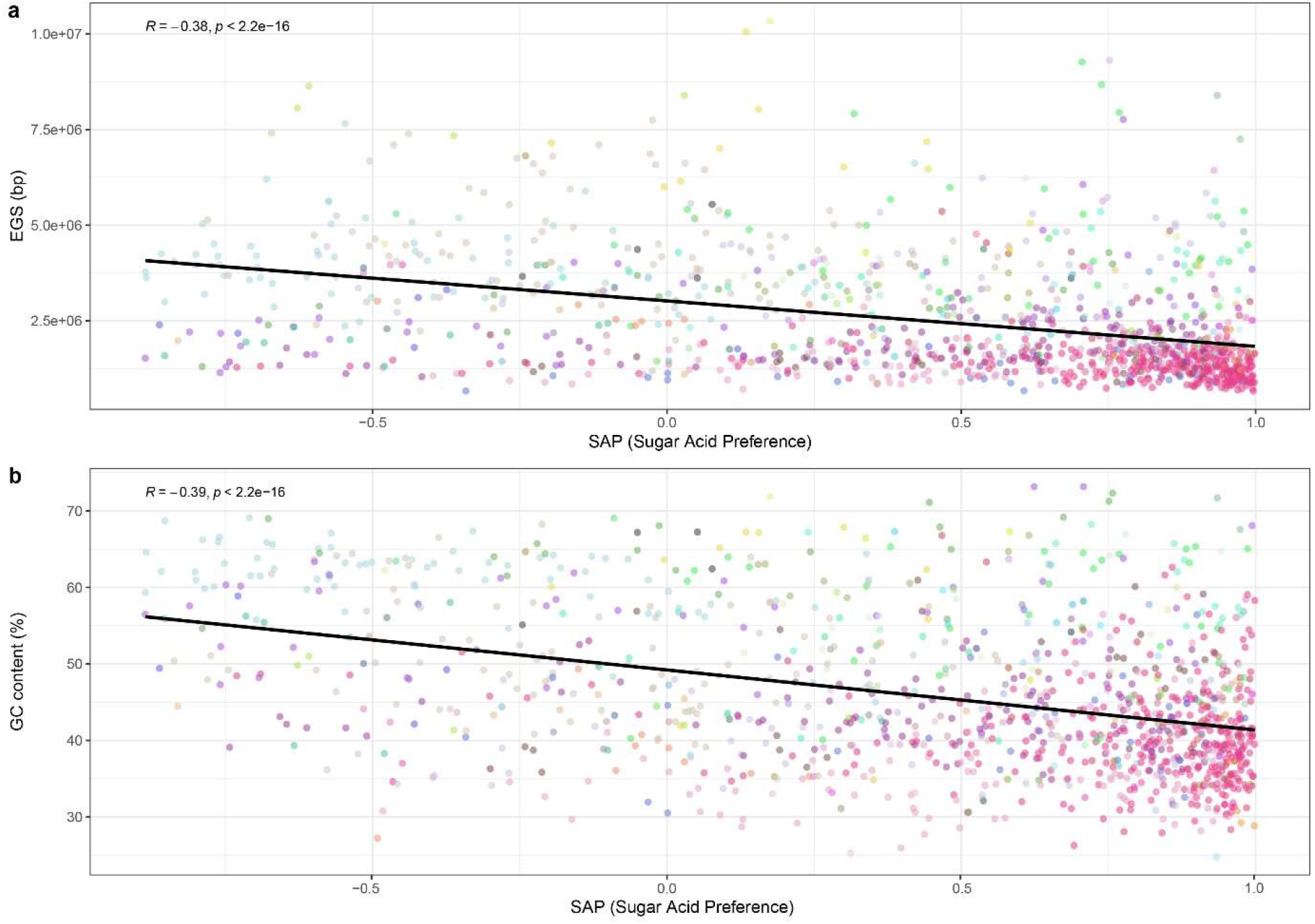
Distribution of calculated SAP (sugar acid preference) index across the range of GC content (a) and estimated genome size (EGS) (b).

**Supplementary Figure S14:**
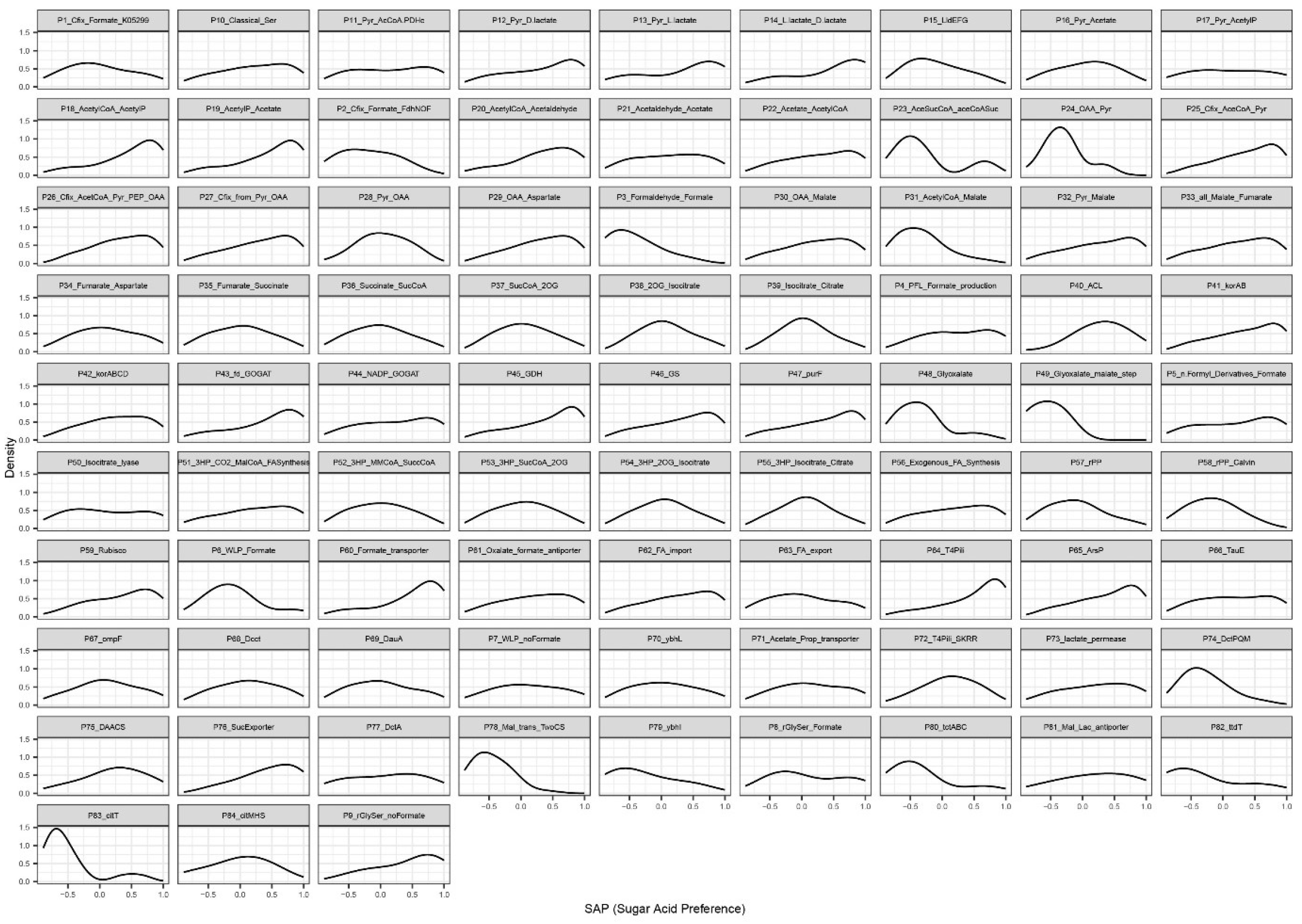
The frequency of MAGs/SAGs encoding genes for 58 modules involved in carbon metabolism as well as genes encoding 25 transporters across the range of sugar acid preference (SAP) index (ranging from 1, indicating sugar specialists, to -1, indicating acid specialists).

**Supplementary Figure S15:**
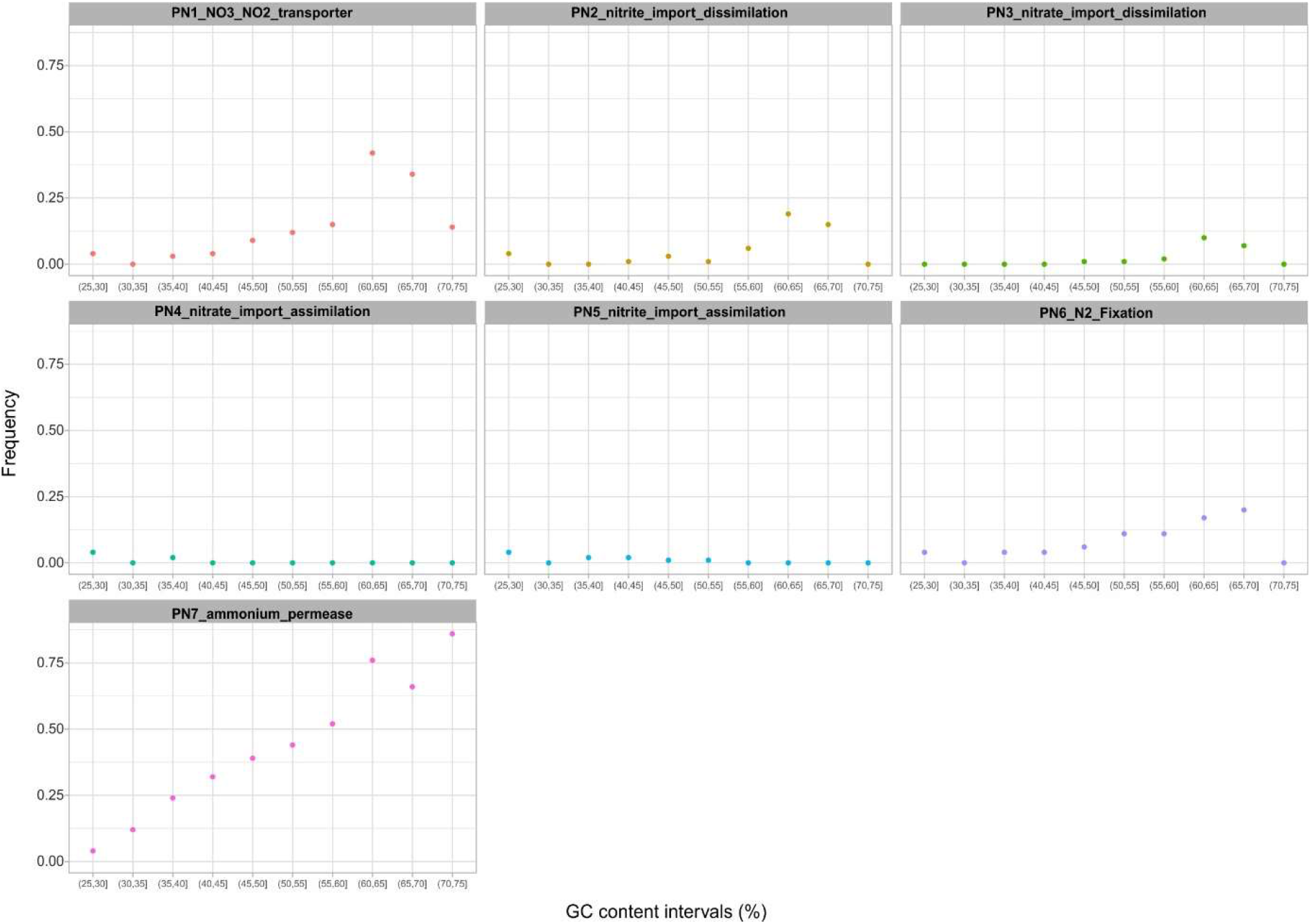
The frequency of MAGs/SAGs containing genes for five modules in nitrogen metabolism as well as genes encoding two transporters across the range of GC content. The number of MAGs/SAGs containing genes for each module is counted in intervals of 5% GC content and normalized by the number of all MAGs/SAGs present in each interval.

**Supplementary Figure S16:**
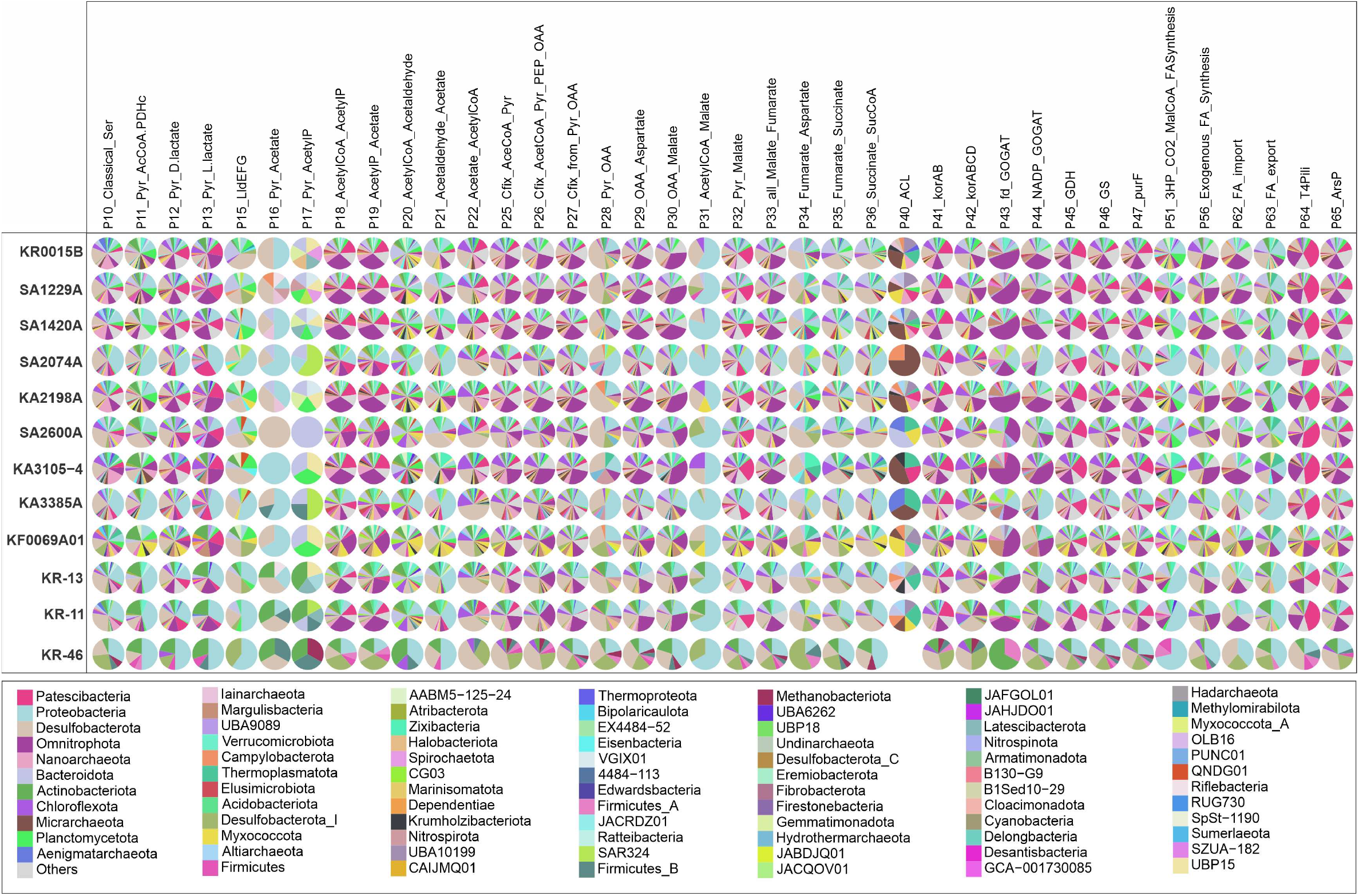
Composition of microbes carrying genes for each metabolic module (other than those in the core) and transporters in different boreholes. The community of microbes encoding each metabolic module in different boreholes is shown as a pie chart. The color legend at the bottom of the plot shows the taxonomic affiliation of MAGs/SAGs encoding different modules at the phylum level. the number of MAGs/SAGs encoding phyla with overall abundance lower than 1% in all boreholes were clumped together as “others” category.

**Supplementary Figure S17:**
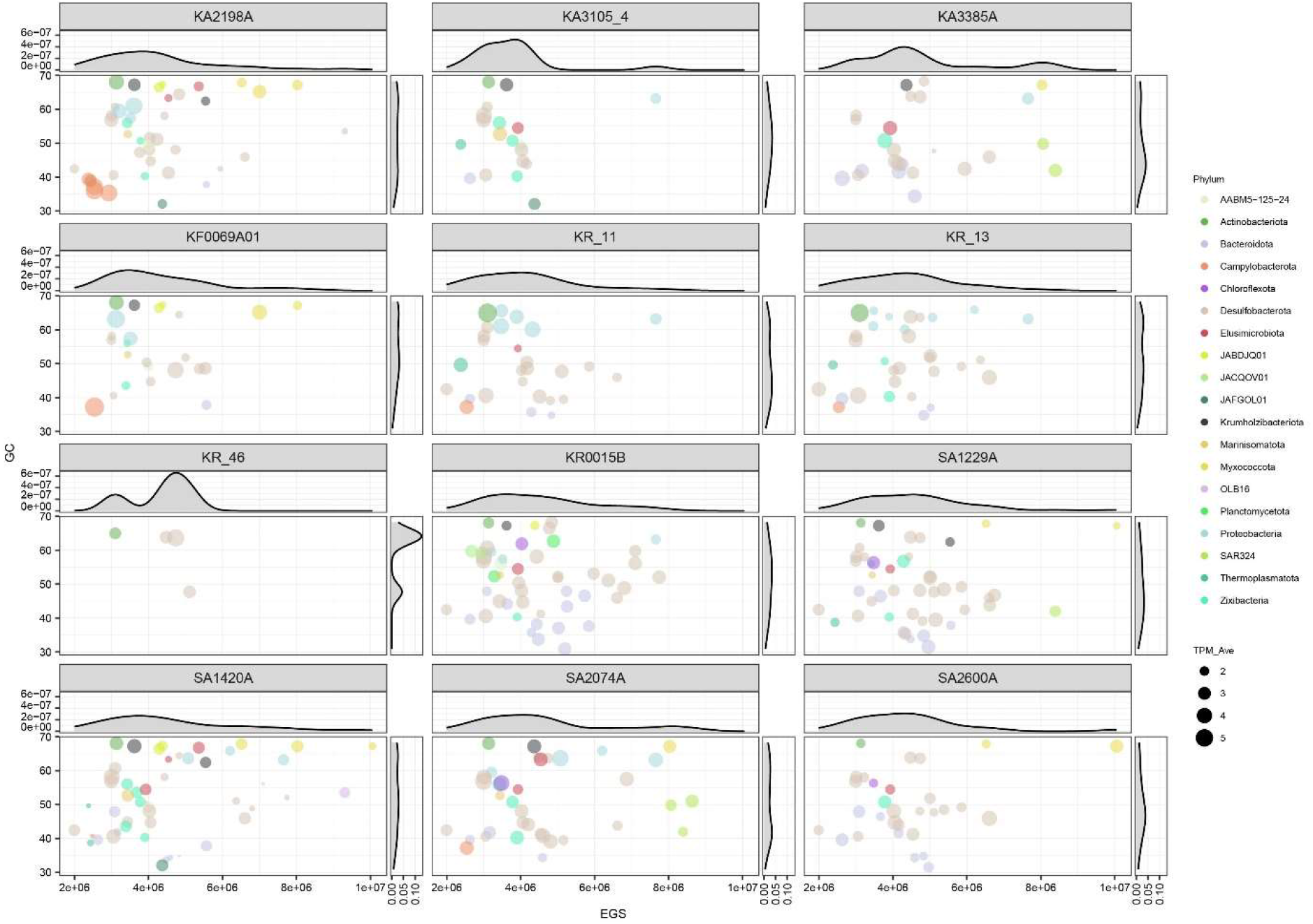
The correlation between GC content and estimated genome size (EGS) of MAGs/SAGs encoding rTCA (encoding at least two of the three key genes) in different boreholes. The color of the circles represents different phyla, while their size indicates the average TPM (calculated as the average of the nonzero log10 TPM values of each rTCA-bearing MAG across all metagenomes sequenced for each borehole).

